# Monoclonal antibodies targeting the rabies virus glycoprotein promotes viral clearance and Fc-dependent neuroprotection in the infected brain

**DOI:** 10.64898/2026.01.13.699195

**Authors:** Seonhee Kim, Anthony Coleon, Florence Larrous, Lauriane Kergoat, Said Mougari, Julien Lannoy, Marta Tiago, David Hardy, Etienne Kornobis, Felix A. Rey, Fabio Benigni, Davide Corti, Hervé Bourhy, Guilherme Dias de Melo

**Affiliations:** Institut Pasteur, Université Paris Cité, Lyssavirus Epidemiology and Neuropathology Unit, F-75015 Paris, France; Institut Pasteur, WHO Collaborating Centre for Reference and Research on Rabies, European Union Public Health Reference Laboratories for Emerging, Rodent-borne and Zoonotic Viral pathogens (EURL-PH-ERZV); Universidade de Lisboa, Faculdade de Medicina Veterinária, 1300-477 Lisboa, Portugal; Institut Pasteur, Université Paris Cité, Histopathology Core Facility, F-75015 Paris, France; Institut Pasteur, Université Paris Cité, Bioinformatics and Biostatistics Hub, F-75015 Paris, France; Institut Pasteur, Université Paris Cité, Plate-forme Technologique Biomics, F-75015 Paris, France; Institut Pasteur, Université Paris Cité, Structural Virology Unit, F-75015 Paris, France; Humabs BioMed SA, a subsidiary of Vir Biotechnology, 6500 Bellinzona, Switzerland

**Keywords:** rabies, therapy, human monoclonal antibody, neutralization, effector functions, animal model

## Abstract

Rabies is a complex disease that has defied efforts to develop an effective treatment. Although it can be successfully prevented through vector control or pre- or post-exposure prophylaxis, no therapeutic options exist for symptomatic rabies, and the disease is still responsible for about 59 000 human deaths each year. Following our successful attempts to use the human monoclonal antibodies RVC20 and RVC58 for intracerebroventricular treatment of rabies, we report here on the effects of these mAbs when administered intravenously and intramuscularly to symptomatic mice. Only a small proportion of mAbs was detected to cross the blood-brain barrier, yet this limited amount sufficiently cleared the viral infection and improved the survival rate of infected mice. Remarkably, these mAbs appeared more effective when used individually than when combined as a cocktail. RVC58 achieved an 80% survival rate, potentially due to its distinct binding topography, potent viral neutralization, and enhanced FcγR engagement in the presence of virions. These findings suggest that monotherapy with a single mAb fully capable of engaging FcγR may be possible, which could substantially lower the costs and improve treatment accessibility in under-resourced settings, potentially contributing to improved equity of global health. Taken together, this study supports mAb therapy as a promising preclinical approach for symptomatic rabies and provides a foundation for further development of an accessible treatment option.

## Introduction

Rabies is a zoonotic viral disease that leads to progressive but lethal encephalitis and is primarily transmitted to humans through the bite of an infected animal, typically a dog (1). The disease is entirely preventable if individuals exposed to the virus receive prompt and appropriate post-exposure prophylaxis (PEP), which consists of the rabies vaccine and rabies immunoglobulins (RIG) derived from human or equine sources (2, 3, 4). An estimated 100,000 individuals receive rabies PEP annually in the United States alone (5), rising to over 5 million in India (6). However, the widespread use of RIG is constrained by factors such as limited availability, high cost, and safety concerns, including the risk of anaphylaxis (7). A study from China showed that out of 10,971 collected rabies cases from 2006 to 2012, only 11.7% of severely wounded individuals (WHO category III bites) initiated PEP vaccination, and a mere 3.9% exposures received RIGs (8), despite the officially recommended regimen for category III cases is to combine vaccination with RIG administration (3). The burden of rabies is disproportionately concentrated in Africa and Asia, particularly in socioeconomically disadvantaged rural areas. Faced with these apparent challenges, several rabies virus (RABV)-specific monoclonal antibodies (mAb) have been developed as potential replacements for human and equine RIGs (7), including Rabishield (9) and Twinrab (10) for the Indian market, among others still in development (7). For individuals whose rabies exposure is not effectively managed, clinical symptoms may emerge within one to three months, though incubation periods can range from less than 10 days to several years (1). The prevailing view is that once RABV reaches the central nervous system (CNS) — prior to the manifestation of clinical symptoms — the disease is invariably fatal (11). Currently, rabies is responsible for an estimated 59,000 deaths annually, with 40% of cases occurring in children under the age of 15 (12). Developing rabies treatments has been notoriously difficult. The first documented human survival of symptomatic rabies was reported in 2004 following application of the Milwaukee protocol (13). Since then, repeated attempts to treat rabies with this protocol have yielded no reproducible benefit (14, 15), and now this approach is discouraged by WHO (1). Other approaches using antivirals have also been proposed, but none of them demonstrated to be potentially successful *in vivo* (16, 17, 18, 19).

As demonstrated for rabies PEP, mAbs may provide a scalable and safe option for rabies therapy. We have previously identified two human mAbs, RVC20 and RVC58, with unique features: i) they bind to two distinct antigenic sites on the RABV glycoprotein (the main surface antigen), ii) they potently neutralize a broad spectrum of RABV isolates *in vitro* and are protective when used as PEP in hamsters, and iii) they exhibit broader range of neutralization across different lyssavirus than other mAbs developed to date (20, 21, 22). Beyond their ability to neutralize virus particles, RVC20 and RVC58 also modulate the immune system via Fc-dependent functions (23). In our previous study, intramuscular administration at the site of infection combined with intracerebroventricular delivery via continuous pump infusion of the RVC20+RVC58 cocktail cleared the virus from the brain and significantly improved clinical conditions in symptomatic rabid mice (24). While these findings are highly encouraging, a treatment against rabies would need to be quickly accessible, affordable, and deliverable through simpler administration routes.

Here we show that the intravenous administration coupled with an intramuscular injection of RVC20+RVC58 mAbs clearly improves survival of symptomatic mice. Following the first injection, these mAbs control virus spread in the brain and modulate the immune response, associated with specific cytokine profiles and microglial activation. Notably, RVC58 administered as monotherapy achieved 80% of survival and clinical recovery, outperforming cocktail treatment in our mice model. Treatment efficacy was time-dependent, with therapeutic outcomes inversely correlated with the interval between symptom onset and mAb administration, underscoring the critical importance of early clinical diagnosis for treatment success. Taken together, these findings suggest that RVC58 represents a promising candidate for rabies therapy, particularly for individuals who fail to receive or experience delays in PEP administration.

## Results

### Intravenous + intramuscular delivery of RVC20+RVC58 mAbs cocktail improves the survival rate of symptomatic rabid mice

To define the onset timepoint of clinical rabies in our Balb/c mouse model, we infected the animals with a 4000 FFU of a field RABV strain in the gastrocnemius muscles, which results in synchronized CNS invasion. With this scheme, RABV was detected in the spinal cord and in the brainstem (composed of diencephalon, midbrain, pons, medulla oblongata) and in the cerebellum as early as 5 days post-infection (dpi), and to a lesser extent also in the telencephalon (constituted by the cerebral cortex, hippocampus, olfactory bulbs) (Figure 1A-C). Clinical signs, including body weight loss and paralysis, were not observed until 8 dpi during standardized clinical scoring, indicating a substantial delay between CNS invasion and symptom onset (Figure 1D).

**Figure 1.**
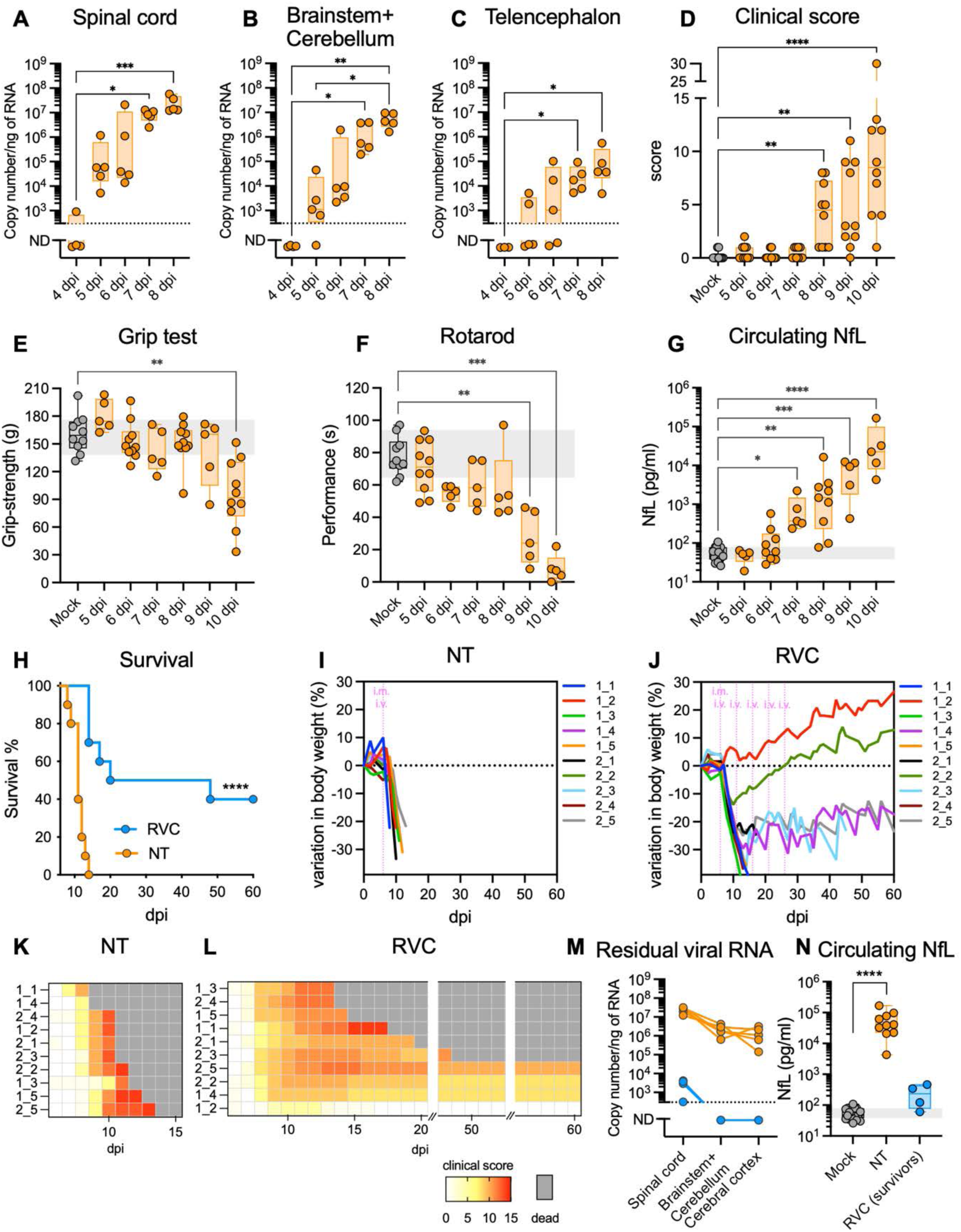
Therapeutic efficacy of intravenous injection of the RVC20 and RVC58 monoclonal antibody cocktail in mice with symptomatic rabies. **A-C.** Rabies virus (RABV) kinetics in the central nervous system of intramuscularly infected mice. Viral load, expressed as copy number/µg of RNA in the spinal cord (A), in the brainstem (composed of diencephalon, midbrain, pons, medulla oblongata) and cerebellum (B) and in the telencephalon (constituted mainly by the cerebral cortex, hippocampus and olfactory bulbs) (n=5/time-point). ND = non detected. **D.** Follow-up of the clinical score in RABV-infected mice at different time-points post-infection (n=10/time-point, 2 independent experiments). **E.** Grip test to measure forelimbs strength in RABV-infected mice at different time-points post-infection (n=10 for mock, 6, 8 and 10 dpi; n=5 for 5, 7 and 9 dpi). **F.** Rotarod test to measure motor coordination in RABV-infected mice at different time-points post-infection (n=10 for mock and 5 dpi; n=5 for 6-10 dpi). **G.** Neurofilament light chain (NfL) levels in the serum of RABV-infected mice at different time-points post-infection (n=19 for mock; n=5 for 5, 7, 9 and 10 dpi; n=9 for 6 and 8 dpi). **H.** Cumulative Kaplan–Meier survival curves of non-treated mice (NT) and mice receiving the RVC20 and RVC58 monoclonal antibody cocktail by IV+IM at 6 dpi. Log-rank (Mantel–Cox) test to compare the groups: **** p<0.0001 (n=10 per group, 2 independent experiments). **I-J.** Body weight progression of NT mice (I) and mice receiving the RVC20 and RVC58 monoclonal antibody cocktail by IV+IM at 6 dpi (J). Mice were weighed daily during the treatment administration and then twice a week up to 60 days post-infection. Each line represents one animal throughout time (n=10 per group, 2 independent experiments). Vertical pink lines indicate the time-points of each injection during the treatment. **K-L.** Individual follow-up of the clinical score of mice under different treatments. Each line represents one animal throughout time (n=10 per group, 2 independent experiments). **M.** Residual viral RNA in the spinal cord, brainstem + cerebellum and cerebral cortex of non-treated animals (NT, orange lines) or of mice that survived after RVC cocktail treatment (60 dpi, blue lines). Each line represents one animal throughout time (n=5 for NT; n=4 for survivors after RVC treatment). **N.** Neurofilament light chain (NfL) levels in the serum of RABV-infected after treatment at 6 dpi (n=19 for mock as also shown in G; n=10 for NT; n=4 for survivors after RVC treatment). Data information: **A-G, N:** Box and whisker plots (median, first and third quartiles, minimum and maximum). Individual values are also shown. Kruskal-Wallis test followed by the Dunn’s multiple comparisons test: asterisks indicate comparisons with the mock-infected group (*p<0.05, **p<0.01, *** p<0.001, **** p<0.0001). The gray crosshatched areas correspond to the 95% CI of the median from the mock-infected mice. Exact P values are shown in the source data tables. **I-L:** The mouse ID is as follows: The first number indicates the replicate, and the second number indicates the mouse within the replicate. For example, 1_3 indicates replicate 1 and mouse 3.

Therefore, to better characterize the symptoms of rabies and refine our model, we monitored animals daily for (i) forelimbs muscular strength using the grip test, (ii) motor coordination on a rotarod apparatus, and (iii) circulating levels of neurofilament light chain (NfL) as a biomarker of neuroaxonal lesion. Weakening of the forelimbs became evident only at 10 dpi (Figure 1E), although the infected mice displayed a gradual impairment of the motor coordination beginning at 6 dpi (Figure 1F). In addition, the serum levels of NfL also showed a gradual increase overtime starting at 6 dpi (Figure 1G). Although no significant changes were observed in body weight or clinical score at 6 dpi, this comprehensive virological, behavioral and serological evaluation indicates that mice already exhibit signs of early symptomatic rabies by 6 dpi.

We next compared the progression of rabies in mice treated with the RVC20+RVC58 human mAb cocktail. The treatment consisted of five intravenous (IV) injections of 25+25 mg/kg of the RVC20 and RVC58 mAbs, one injection per day every five days, starting at 6 dpi. The animals also received a single intramuscular (IM) injection at a lower dose of 2+2 mg/kg of the mAb cocktail at 6 dpi to quickly limit further viral replication at the injection site. Mock-infected animals received the same RVC mAbs treatment regimen, while non-treated animals received saline only. From this experiment, we observed that the IV+IM treatment with RVC20+RVC58 resulted in a survival rate of 40% (4 out of 10) of animals up to 60 dpi (Figure 1H; p<0.0001).

The non-treated mice showed rapid weight loss accompanied by progressive development of neurological symptoms (Figure 1I,K) and eventually reaching the humane endpoint between 8 and 14 dpi. The mice treated at 6 dpi with the RVC cocktail initially exhibited serious decline in weight and movement from 8 dpi, similar to non-treated animals (Figure 1J). However, the weight loss slowed following the second IV injection at 11 dpi. Despite the early development of monoplegia in 20% of treated animals (2 out of 10 mice at 8 dpi), symptoms were maintained or resolved over time in the treated group (Figure 1L). By the end of the experiment at 60 dpi, very low or no viral RNA was detected in the CNS of the surviving animals (Figure 1M), and NfL levels in their serum were as low as in mock-infected mice (Figure 1N). Clinical scores differed significantly between non-treated and RVC-treated animals (one-way ANOVA, p<0.001) with statistical power exceeding 0.9 at a sample size of 10 per group.

We additionally tested treatments initiated at 7 or 8 dpi to assess the effect of delayed intervention. One or two days of delay in treatment dramatically lowered the survival rate of animals: 20% (2 out of 10) of animals treated at 7 dpi survived, with an average lifespan of 17.8 days after the first RVC injection. No animal survived when treatment began at 8 dpi. Overall, the intramuscular and intravenous administration of RVC20+RVC58 mAb cocktail improved the survival rate of infected animals and delayed disease progression, with inversely correlated efficacy to the time delay between onset of early symptoms and treatment initiation.

### mAb cocktail treatment via IV + IM routes limits virus replication in the CNS despite a low uptake of antibodies across the blood-brain barrier

To understand how the RVC cocktail treatment via the IV + IM routes rescues animals from early symptomatic rabies, mice received a single high-dose IV injection of the RVC20+RVC58 mAb cocktail (25+25mg/kg) combined with a low-dose IM injection (2+2 mg/kg) at dpi 6. Samples were collected at defined time points (Figure 2A). To investigate the contribution of Fc-mediated effector functions to the treatment, we also tested the LALA variants of these mAbs (RVC20-LALA and RVC58-LALA; hereafter called LALA). The LALA mAbs carry two point-mutations on their Fc region (L234A and L235A) that abolish their binding affinity to Fcγ receptors (FcγRs) and complement.

**Figure 2.**
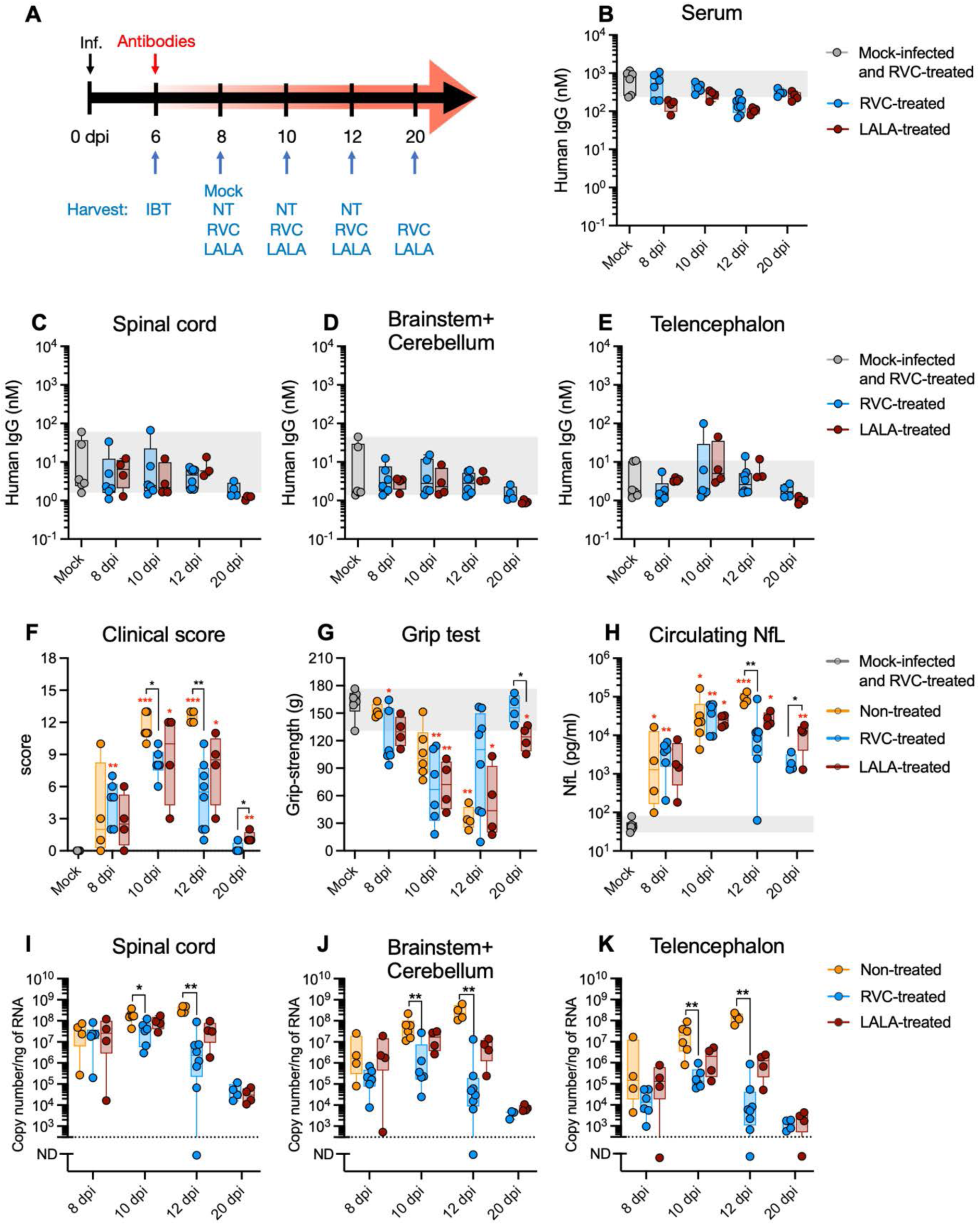
Time-dependent activity of the RVC20 and RVC58 monoclonal antibody cocktail injected intravenously in mice with symptomatic rabies. **A.** Timeline of the experiment. The animals were treated at 6 days post-infection (dpi) and samples were collected at different time-points. IBT = infected, before treatment. Mock = mock-infected and RVC-treated animals. NT = infected, non-treated. RVC = infected, RVC20+RVC58 treated animals. LALA = infected, LALA20+LALA58 treated animals. **B-E.** Concentration of human IgG in the serum (B), in the spinal cord (C), in the brainstem + cerebellum (D) and in the telencephalon (E). **F.** Follow-up of the clinical score. **G.** Grip test to measure forelimbs strength. **H.** Neurofilament light chain (NfL) levels in the serum. **I-K.** Rabies virus load in the spinal cord (I), in the brainstem + cerebellum (J) and in the telencephalon (K). Data information: Box and whisker plots (median, first and third quartiles, minimum and maximum). Individual values are also shown. Mock (n=6); Non-treated (n=4 for 8 dpi; n=6 for 10 dpi; n=4 for 12 dpi); RVC (n=6 for 8 and 10 dpi; n=8 for 12 dpi; n=4 for 20 dpi); LALA (n=4/time-point). 2 independent experiments. Kruskal-Wallis test followed by the Dunn’s multiple comparisons test: red asterisks indicate comparisons with the mock-infected group, whereas black asterisks indicate comparisons among treatments. Unpaired T test to compare RVC and LALA at 20 dpi (*p<0.05, **p<0.01, *** p<0.001, **** p<0.0001). The gray crosshatched areas correspond to the 95% CI of the median from the mock-infected mice. Exact P values are shown in the source data tables.

First, we performed an ELISA test to verify if the injected antibodies could be detected in the serum and in different regions of the CNS at 2, 4, 6 or 14 days after the single IV + IM administration, which corresponds to 8, 10, 12 or 20 dpi (Figure 2B-E). Overall, the CNS exhibited approximately 1000-fold lower concentrations of antibodies in mass fraction compared to the serum. RVC and LALA cocktail treatments did not show differences in antibody uptake into the CNS, and the level of uptake remained consistent across all CNS regions and across different time points.

Prior to harvest, the animals were monitored daily for clinical signs and behavior, which were systematically scored to track changes in their health status (Figure 2F-G). Despite the relatively low levels of antibody uptake, both RVC- and LALA-treated animals showed a slower progression of symptoms compared to non-treated controls. By 10 dpi, the symptoms of treated mice notably improved as they were more alert and active than the previous time-point. At 20 dpi, the clinical scores of treated animals were lower than those recorded at 8 dpi, indicating a gradual recovery over time (Figure 2F). Additionally, while all animals initially exhibited a similar decline in forelimb muscle strength, RVC and LALA-treated animals began to regain strength around 12 dpi and continued to recover thereafter, in contrast to non-treated mice. Furthermore, RVC-treated mice displayed faster recovery in strength with a significant difference compared to LALA-treated animals at 20 dpi (Figure 2G). These physical manifestations of recovery under antibody treatment were also supported by physiological evidence. We measured the circulating NfL in collected serum samples, which revealed significantly lower concentration in RVC-treated animals compared to non-treated counterparts at 12 dpi (Figure 2H). By 20 dpi, NfL levels were lower in RVC-treated animals than in LALA-treated mice.

To evaluate the antiviral efficacy of antibody treatment, we quantified viral RNA levels across different CNS regions (Figure 2I-K). High viral loads were detected in the majority of spinal cords and brains at 8 dpi. Beyond this time-point, viral RNA levels in all tissues of non-treated animals continued to rise, whereas RVC-treated animals showed a progressive decline relative to non-treated group, with a significant reduction by 10 dpi in the spinal cord, in the brainstem + cerebellum, and in the telencephalon. In these regions, the viral RNA levels were then low or undetectable at 12 dpi and 20 dpi (Figure 2I-K).

Taken together, our findings indicate that a combination of a single IV + IM administration of the RVC20+RVC58 cocktail at 6 dpi reaches the CNS in limited quantities but is sufficient to reduce the virus replication and spread. This result is consistent with the observed improvement of the clinical symptoms and the reduction in neuronal damage, which may collectively contribute to improved survival rates. Furthermore, intact interaction with FcγRs appears important for the full therapeutic efficacy of the RVC cocktail against RABV infection, as shown that RVC-treated mice exhibited better clinical recovery and earlier reduction of the CNS viral load compared to LALA-treated mice.

### mAb administration modulates the immune environment within the CNS

To characterize the CNS immune environment in RVC20+RVC58 mAb-treated animals, we evaluated the expression of immune mediator genes across different CNS regions at 2, 4, 6 or 14 days after treatment (corresponding to 8, 10, 12 or 20 dpi). Incorporating all time-points and CNS regions together, different groups of mediators contributed to the distinct clustering of NT and RVC animals in a principal component analysis. Whereas *Ifnb* and *Il6* were most related to NT, *Ifng* and *Tgfb* mostly associated with the RVC-treated group. This observation suggests that the balance between pro- and anti-inflammatory mediators plays an important role in creating an effective antiviral environment against RABV infection (Appendix Figure S1A-B).

The most pronounced gene expression change was observed for the pro-inflammatory cytokines *Ifnb* and *Il6*, which were significantly elevated as early as 8 dpi in non-treated animals. Their expression in RVC-treated mice was relatively limited, with significant reduction compared to NT at 10 dpi (Figure 3A-B). The differences of other tested pro-inflammatory cytokines, *Ifng* and *Tnfa*, were less evident, however, their expression tended to be lower in RVC than in LALA animals (Appendix Figure S1C-D). Regarding anti-inflammatory cytokines, the gene expression of *Tgfb* was higher in the spinal cords of RVC-treated animals than in non-treated animals at 10 dpi, whereas *Il10* expression was elevated throughout the CNS, with a diminution in RVC-treated animals at 20 dpi (Appendix Figure S1E-F). Finally, the expression of the interferon-related gene *Ifit2* was significantly lower in the CNS of RVC-treated animals between 10 and 12 dpi (Appendix Figure S1G).

**Figure 3.**
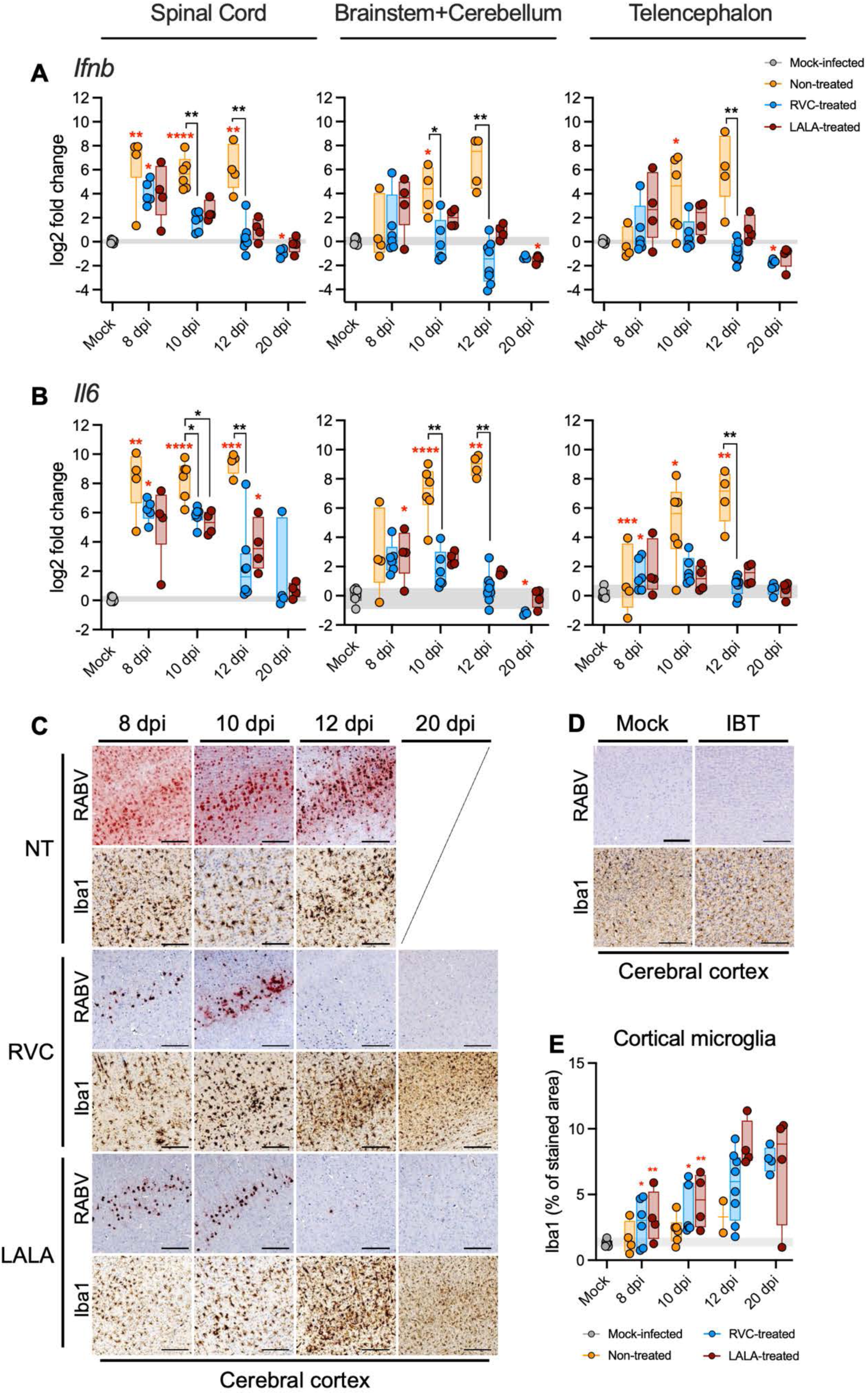
Modulated immune response in the brain under RVC20 and RVC58 monoclonal antibody cocktail treatment. **A-B.** Time-dependent gene expression of interferon beta (A) and interleukin 6 (B) in the spinal cord, brainstem + cerebellum, and telencephalon of mice under different treatments. C-D. Immunohistochemical analysis of the cerebral cortex showing staining for the rabies virus (RABV, red spots) and for microglia (Iba1, brown cells) in mice under different treatments at different days post-infection. Representative images. Scale bar = 200 µm. NT = infected, non-treated. RVC = infected, RVC20+RVC58 treated animals. LALA = infected, LALA20+LALA58 treated animals. Mock = mock-infected and RVC-treated animals. IBT = infected, before treatment. E. Quantification of cortical microglia in the brain of mice under different treatments at different days post-infection. Data information: Box and whisker plots (median, first and third quartiles, minimum and maximum). Individual values are also shown. Mock (n=6); Non-treated (n=4 for 8 dpi; n=6 for 10 dpi; n=4 for 12 dpi); RVC (n=6 for 8 and 10 dpi; n=8 for 12 dpi; n=4 for 20 dpi); LALA (n=4/time-point). 2 independent experiments. Kruskal-Wallis test followed by the Dunn’s multiple comparisons test: red asterisks indicate comparisons with the mock-infected group, whereas black asterisks indicate comparisons among treatments. Unpaired T test to compare RVC and LALA at 20 dpi (*p<0.05, **p<0.01, *** p<0.001, **** p<0.0001). The gray crosshatched areas correspond to the 95% CI of the median from the mock-infected mice. Exact P values are shown in the source data tables.

An immunohistochemical study to stain RABV and microglia corroborated the above-mentioned findings. While a diffuse staining of RABV was detected throughout the brain as early as 8 dpi in NT, RVC- and LALA-treated animals presented mild RABV staining restricted to 8 and 10 dpi (Figure 3C and Appendix Figure S2). Microglia responded to the treatment despite the decreasing viral load, presenting a progressive increase in the Iba1 staining in RVC- and LALA-treated animals compared to NT or mock-infected animals (Figure 3C-E and Appendix Figure S2). Given that *Ifnb* and *Il6* are mainly derived from microglia in the CNS (25), we sought to specifically characterize the effects of the RVC treatment on microglia. For this purpose, we isolated microglia from whole brains of mice at 9 dpi (Figure 4A) and subjected it to bulk RNA sequencing, comparing sorted microglia in NT infected mice, and mice treated with RVC or LALA cocktails (one IV+IM administration at 6 dpi), with mock-infected animals serving as reference.

**Figure 4.**
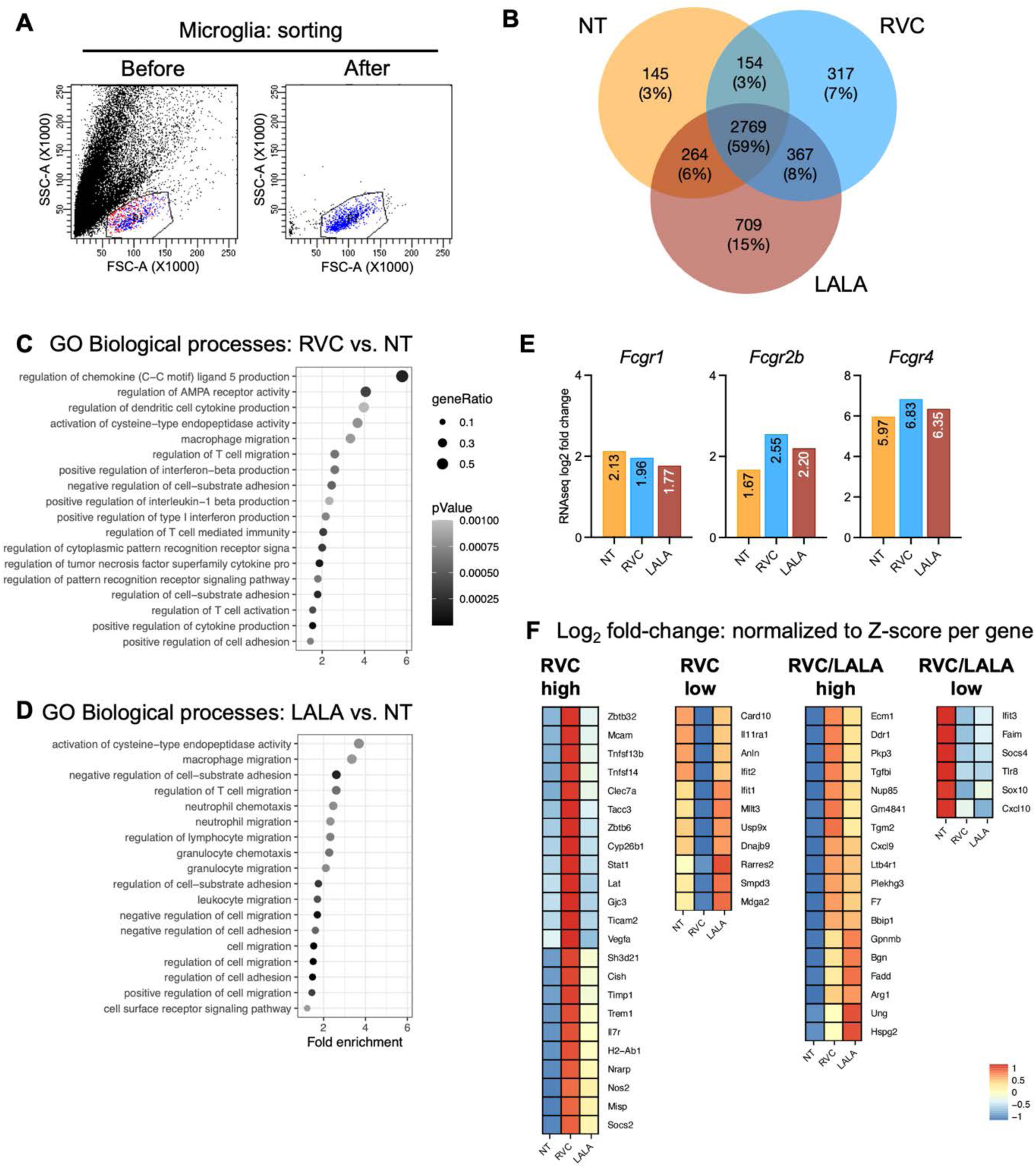
Purified microglia show differential expression of immunity-associated genes under RVC20 and RVC58 monoclonal antibody cocktail treatment. **A.** Representative dot plot showing microglial cells isolation before and after cell sorting. **B.** Venn diagram of microglial cells RNAseq showing the unique and overlapping differentially expressed genes (DEGs) of non-treated (NT), RVC-treated and LALA-treated animals. The number of genes is shown in the diagram. Each condition was normalized with the mock-infected animals (n=3/group). **C-D.** GO enrichment analysis considering biological process of RVC in comparison with NT (C), and of LALA in comparison with NT (D). Circle sizes are proportional to the gene ratio, which shows the total size of the gene set associated with the GO terms. Circle color is proportional to the corrected p values. **E.** Differential expression of the Fc gamma receptors *Fcgr1*, *Fcgr2b* and *Fcgr4* in microglial cells from the brain of mice under different treatments. The log2 fold change values in comparison with the mock-infected animals are shown in the graphs (n=3/group). **F.** Heatmap showing the differential expression of selected immune-related genes in microglial cells from the mice under different treatments. The expression of individual genes from the three groups was centered and standardized to Z-score, with 59 genes presenting differences bigger than 1.5 log2 fold-change between RVC and NT groups. The color gradient represents the Z-score of each gene compared with the mock-infected animals (n=3/group).

Compared to the mock-infected group, microglia of non-treated infected animals presented 3,332 differentially expressed genes (DEGs; increased or decreased) while RVC-treated showed 3,607 DEGs, and LALA-treated animals presented 4,109 DEGs. Among these, 2,769 DEGs were common to all three groups (Figure 4B) and were subjected to further analysis, focusing on genes associated with immune response-related Gene Ontology (GO) terms in the Biological Process category. In comparison to NT animals, the DEGs from RVC-treated animals were mostly associated with cell-mediated immunity and innate immunity (Figure 4C), whereas the DEGs of the microglia from LALA-treated animals were mainly associated with cell migration and adhesion (Figure 4D).

Of particular interest with respect to effector function was the expression of FcγRs by microglia in the brain of rabid animals. We detected that RABV infection induced the expression of both activating (*Fcgr1* and *Fcgr4*) and inhibitory (*Fcgr2b*) FcγRs on microglial cells (Figure 4E). While the expression of *Fcgr1* was not affected by the mAbs treatment, microglia from RVC-treated animals exhibited higher expression of both *Fcgr2b* (increase of 52.1%) and *Fcgr4* (increase of 14.4%) compared to NT. This increase was also observed in microglial cells from LALA-treated animals, but to a lesser extent.

We next examined the DEGs among the different groups to identify molecular signatures associated with therapeutic efficacy. Among the 2,769 DEGs common to all three groups, 791 genes were related to immune response (Figure 4F). Among the DEGs with notable differences (absolute fold change between RVC and NT > 1.5), we detected four patterns of expression: 23 DEGs with high expression in RVC compared to NT and LALA (named RVC high), 11 DEGs with elevated expression in NT and LALA but not in RVC (named RVC low), 18 DEGs showed high expression in both RVC and LALA compared to NT (RVC/LALA high), and 6 DEGs were elevated in NT but not in RVC nor in LALA (named RVC/LALA low). Several genes in the RVC high category have neuroprotective roles, including *Tnfsf13b* (BAFF), *Tnfsf14* (LIGHT), *Timp1*, and *Vegfa*. Furthermore, among the RVC low, some genes are related to apoptotic activities or are involved in brain damage, such as *Card10, Rarres2* (Chemerin)*, Smpd3* and *Mdga2*. All these findings support a neuroprotective effect of the RVC treatment. Finally, among the DEGs increased in both RVC and LALA, we detected some targets classically related to microglia polarization, both M1 (*Cxcl9, Nos2, Ecm1*) and M2 (*Arg1, Vegfa, Tgm2*).

In light of these findings, we also evaluated the T and B lymphocyte populations in the brain at 12 dpi. We chose this late time-point that is close to the humane endpoint for the NT animals, since RABV infection does not result in strong lymphocyte infiltration in the brain. Indeed, very low numbers of T cells were observed in the brains of NT mice at 12 dpi (Figures EV1-2), however, IV+IM treatment with mAbs altered this outcome. Both the RVC and LALA treatments increased the population of T lymphocytes in the cerebral cortex and brainstem, particularly in the pons and in the medulla oblongata (Figure EV1). The population of B lymphocytes were very limited in all groups, and almost absent in the cerebellum of RVC animals (Figure EV2). Collectively, these findings suggest that the IV+IM mAb treatment promotes a modest but notable recruitment of T lymphocytes into the CNS, as anticipated in ontology analysis, pointing to a T cell-mediated contribution to the therapeutic effect.

### Distinct spatial arrangements with RABV may drive differences in the performance of RVC20 and RVC58

We first verified the neutralization potency of RVC20, RVC58 or the RVC20+RVC58 cocktail against four lyssaviruses: the *Lyssavirus rabies* (RABV) strain used to infect the animals in this study (isolate 8743THA), the cell culture-adapted rabies challenge virus (CVS-11), a *Lyssavirus rabies* adapted to vampire bat (isolate 9001FRA), and the *Lyssavirus hamburg* (EBLV-1, isolate 8918FRA). As expected, both RVC20 and RVC58 potently neutralized all the tested viruses (IC50 < 10 ng/mL). However, the IC50 values of RVC58 were 3 to 12-fold lower than those of RVC20, depending on the RABV strain (Figure 5A), in agreement with previous data (20).

**Figure 5.**
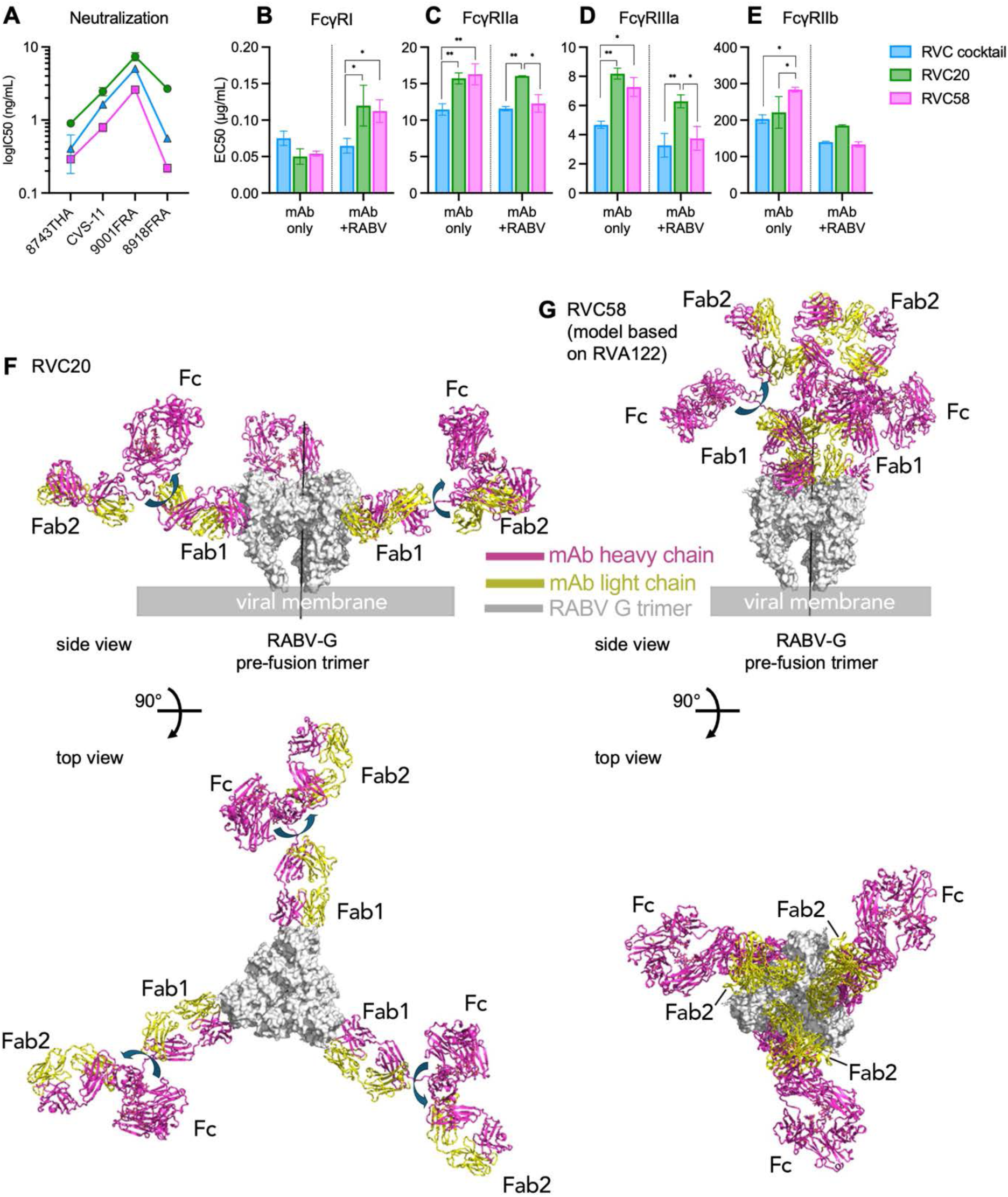
RVC20 and RVC58 mAbs approach differently to the rabies virus glycoprotein (RABV-G). **A.** Neutralization of different Lyssaviruses with the RVC cocktail (1:1 mixture of RVC20 + RVC58) or with RVC20 or RVC58 alone on BSR cells. The logIC50 values are expressed as mean ± SD (n=2 independent experiments). **B-E.** RVC20 and RVC58 binding to human FcγRI (B), FcγRIIa (C), FcγRIIIa (D), and FcγRIIb (E) as single mAbs or in cocktail, alone or pre-incubated RABV. The IC50 values were obtained after fitting the data to a four-parameter regression equation with 1/y^2^ weighting and are expressed as mean ± SD (n=2 independent experiments). Two-way ANOVA test followed by the Tukey’s multiple comparisons test: only comparisons between RVC20 and RVC58 are shown in the graphs (ns: non-significant, *p<0.05, **p<0.01). Exact P values are shown in the source data tables. **F-G.** Modelling of the full-length mAbs RVC20 (F) and RVC58 (G) bound to the pre-fusion trimer of the RABV-G, generated as described in Materials and Methods. RABV-G is shown in gray surface representation with the 3-fold molecular axis drown as a vertical black line at the center, and the antibodies as ribbons with the heavy and light chains colored as indicated. Only one of the two Fabs can bind to a given G trimer (labeled Fab1), with the second one (Fab2) potentially binding an adjacent G trimer. Note that the hinge region between Fab and Fc is highly flexible (illustrated by the curved arrows), and only Fab1 is rigidly maintained against the G trimer, projecting Fab2 and the Fc region away from the bound G trimer differently. For RVC20, Fab 1 projects perpendicular to the G trimer axis (in a plane parallel to the viral membrane), while for RVC58 it projects vertically, away from the viral membrane. The range of distances the Fc samples to reach an Fcγ receptor of an effector cell is different for the two mAbs. The Figure also shows that the epitope of RVC20 may not be accessible on G trimers densely packed at the viral surface, while that of RVC58 would be exposed in all G trimers. RVC20 could also interfere with RABV-G trimers clustering at the infected cell surface for budding of new particles, contrary to RVC58, highlighting the different potential modes of action of the two antibodies.

To evaluate Fc-mediated effector functions, we assessed the binding capacity of RVC20, RVC58, or the RVC20+RVC58 cocktail to the human activating receptors FcγRI, FcγRIIa, FcγRIIIa (orthologous to murine FcγRIV), and to the human inhibitory receptor FcγRIIb, in the absence or presence of RABV (isolate 8743THA). For virus-bound conditions, we incubated a constant amount of RABV with varying concentrations of the mAbs, as in a neutralization assay, and then measured the binding of the mAb-RABV complexes to the FcγRs by luminescence (Figure 5B–E). In the absence of RABV, both RVC20 and RVC58 bound similarly to all activating FcγRs, while the RVC cocktail showed lower EC50 values for FcγRIIa and FcγRIIIa under these conditions. In the presence of RABV, the EC50 values of the mAbs generally decreased for all FcγRs, except for FcγRI. Notably, the EC50 values of RVC58 binding to FcγRIIa and FcγRIIIa decreased considerably compared to RVC20 (Figure 5C–D).

To better understand the functional differences between RVC20 and RVC58, we modeled their binding to the pre-fusion trimer of the RABV glycoprotein (RABV-G). Based on the experimentally determined structure (21, 22), RVC20 binds to RABV-G laterally (antigenic site I) (22, 23), positioning its Fc regions in a plane parallel to the viral membrane (Figure 5F, Movie EV1). In contrast, RVC58 binds to RABV-G at the G-trimer apex (antigenic site III) and perpendicular to the viral membrane (Figure 5G, Movie EV2). This binding mode potentially results in denser arrangement of Fc regions, which could project further away from the virion’s surface (Figure 5G, Movie EV2). This distinct binding topography may account for enhanced FcγR engagement of RVC58 in the presence of RABV virions, possibly enabling more efficient clearance by FcγR-expressing cells. In contrast, previous studies have shown that RVC20 activates cell-mediated effector functions more potently than RVC58 on RABV-infected cells (23), likely due to the lower density of glycoproteins on the cell surface compared to virions. The lateral binding of RVC20 may therefore favor blockade of virus morphogenesis and be more efficient at killing or phagocytosis of infected cells, highlighting the complementary but distinct modes of action of the two antibodies.

### RVC58 monotherapy improves survival rate in symptomatic rabies

To further unravel the action mechanisms underlying the therapeutic activity of these mAbs, and given the differences between RVC20 and RVC58 in terms of Fab and Fc properties, we conducted additional experiments using single antibodies (RVC20, RVC58, LALA20 or LALA58) in the place of the cocktail. First, we aimed to assess these single antibodies in a “post-exposure prophylaxis” setting, in which the animals were infected with RABV and received a single intramuscular injection of each mAb (20 mg/kg) at 2 dpi. All animals treated with RVC20, RVC58 and LALA58 survived, however, LALA20 protected only 60% (3 out of 5) of the animals (Figure 6A). These results indicate that even at the time point as early as 2 dpi (before viral CNS invasion), the protective efficacy of RVC20 relies heavily on Fc-mediated effector functions, whereas RVC58 primarily retains full protective activity through its Fab-mediated neutralization. By the end of the experiment at 60 dpi, all clinical parameters were normal, but NfL levels were slightly elevated in some animals treated with RVC20 and LALA20 (Figure 6B).

**Figure 6.**
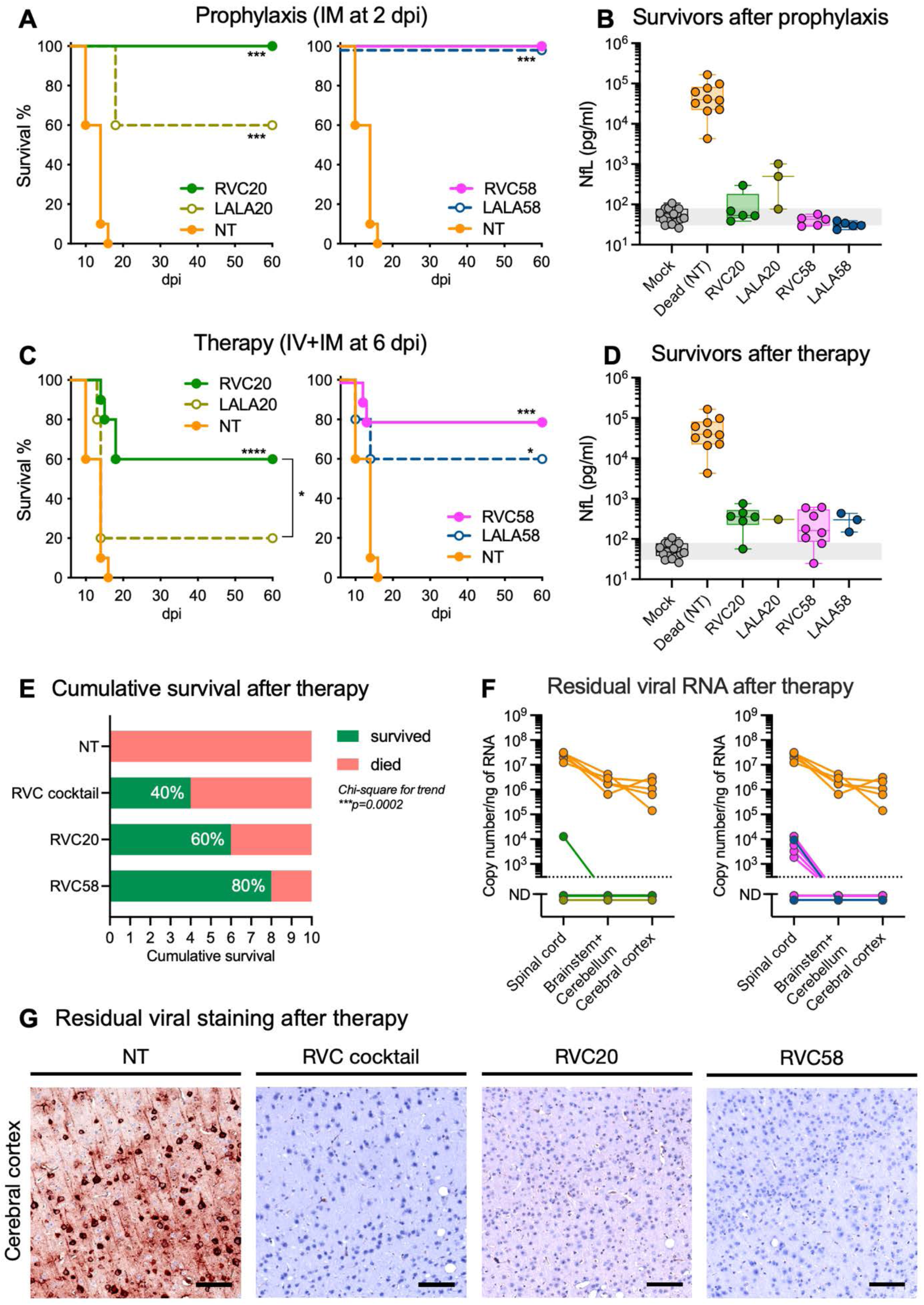
Therapeutic efficacy of intravenous injection of single RVC20 or RVC58 monoclonal antibodies in mice with rabies. **A.** Cumulative Kaplan–Meier survival curves of non-treated mice (NT) and mice treated intramuscularly with a single injection of RVC20, LALA20, RVC58 or LALA58 monoclonal antibodies at 2 dpi to mimic post-exposure prophylaxis. Log-rank (Mantel–Cox) test to compare the groups: *** **p**<0.001 (n=5/group). **B.** Neurofilament light chain (NfL) levels in the serum of RABV-infected surviving animals (collected at 60 dpi) after prophylaxis at 2 dpi. Box and whisker plots (median, first and third quartiles, minimum and maximum). Individual values are also shown. The gray zone corresponds to the 95% CI of the median from the mock-infected mice. Mock (n=19 as also shown in Figure 1G); NT (n=10, dead); RVC20 (n=5), LALA20 (n=3), RVC58 (n=5), LALA58 (n=5). **C.** Cumulative Kaplan–Meier survival curves of non-treated mice (NT) and mice receiving the RVC20, LALA20, RVC58 or LALA58 monoclonal antibodies by IV+IM starting at 6 dpi. Log-rank (Mantel–Cox) test to compare the treated groups with the non-treated animals: *p<0.05, **p<0.01, *** p<0.001 (n=5 for NT, LALA20 and LALA58; n=10 for RVC20 and RVC58, 2 independent experiments). **D.** Neurofilament light chain (NfL) levels in the serum of RABV-infected mice that survived after treatment (60 dpi). Box and whisker plots (median, first and third quartiles, minimum and maximum). Individual values are also shown. The gray zone corresponds to the 95% CI of the median from the mock-infected mice. Mock (n=19 as also shown in Figure 1G); NT (n=10, dead); RVC20 (n=6), LALA20 (n=1), RVC58 (n=8), LALA58 (n=3). **E.** Cumulative survival after intravenous therapy with the TVC20+RVC58 cocktail or the RVC20 and RVC58 mAbs only. Fisher’s exact test for contingency. **F.** Residual viral RNA in the spinal cord, brainstem + cerebellum and cerebral cortex of non-treated animals, or of mice that survived after treatment (60 dpi). **G.** Immunohistochemical analysis showing staining for the rabies virus (RABV, red spots) in the cerebral cortex of NT mice that died, or of mice that survived after treatment (60 dpi). Representative images. Scale bar = 50 µm. Data information: **A, C:** Exact P values are shown in the source data tables. G: These are high-magnification views of the submacroscopic brain images showed in Figure EV5.

Next, we evaluated the same single antibodies in a “therapy” setting, in which the mAbs were administered by IV (25 mg/kg) once every five days starting at 6 dpi, alongside a single low dose IM injection (2 mg/kg) at 6 dpi. At 12 dpi, the viral load and host gene expression profiles induced by the RVC20 or RVC58 monotherapies did not statistically differ from those induced by treatment with the RVC20+RVC58 cocktail (Figure EV3). However, examination of lymphocyte infiltration revealed modest differences in T cell profiles between RVC20 and RVC58 groups, with RVC58 monotherapy yielding T cell levels more comparable to those of RVC cocktail group in the cerebral cortex and pons/medulla oblongata, which are among the first brain regions invaded by RABV in our animal model (Figure EV3J-N).

As previously observed from the RVC20+RVC58 cocktail and LALA counterparts, the efficacy of single mAb monotherapy appeared to depend on both neutralization and effector functions, with RVC20 and RVC58 outperforming their LALA variants (Figure 6C and Figure EV4). By 60 dpi, most clinical parameters were normal, but some animals did not completely recover their body weight (Figure EV4), and NfL levels were still slightly elevated in some surviving animals (Figure 6D). The survival rate of the RVC20 treatment was 60% (6 out of 10), while RVC58 treatment yielded 80% survival (8 out of 10), exceeding both the RVC20 monotherapy and the RVC20+RVC58 cocktail (Figure 6E). Regardless of treatment, viral RNA was undetectable in the brains of survivors but still present at very low levels in some spinal cords (Figures 1M and 6E). Viral isolation assays from qPCR-positive spinal cords were negative (Appendix Figure S3). Additionally, viral staining was negative throughout the entire CNS (Figure 6G and EV5). These results indicate full viral clearance and suggest that RVC58 monotherapy may represent an effective treatment strategy for symptomatic rabies, potentially attributable to potent viral neutralization, distinct binding topography, and enhanced FcγR engagement in the presence of virions.

## Discussion

Rabies is a highly complex disease that has challenged efforts to develop efficient treatments, for which the therapeutic efficacy is highly dependent on the timing of intervention. Although it can be successfully prevented through vector control or pre- or post-exposure prophylaxis and has been largely eliminated in high-income regions, access to effective measures is still limited in low- and middle-income countries, where the burden of disease is greatest (1). Despite being a preventable disease, the high cost and limited accessibility of therapeutic options force rabies to become untreatable and cause otherwise avoidable deaths.

Here, we evaluated the therapeutic potential of the RVC20+RVC58 human mAbs cocktail when primarily injected via IV+IM routes in early symptomatic rabid mice. As expected, only a low amount of mAbs reached the brain parenchyma, consistent with previous studies showing that only 0.1 to 1% of unmodified circulating antibodies accumulate in the brain (26, 27, 28, 29). However, this limited CNS delivery was associated with successful viral clearance and improved survival rate in the present study. Additionally, as we established in our previous study, the administration of a low dose of mAbs at the site of infection is pivotal for local viral clearance and higher treatment efficacy (24). This is why IV administration was combined with a lower dose IM injection in all treatment protocols starting at 6 dpi or later, accounting that mAb accumulation in muscle also occurs at relatively low rates after an IV injection, approximately 4% of the plasma levels (29).

The effectiveness of an anti-rabies therapy should be evaluated not only based on survival rate, viral clearance, but also on quality of life following recovery (14, 30). The RVC20+RVC58 cocktail treatment indeed promoted survival, controlled viral spread, and cleared RABV from the infected mouse brains by 60 dpi. From around 15 dpi onward, mAb-treated mice entered a recovery phase, in which clinical scores and muscular strength, as measured by grip test, stabilized and eventually returned to near normal levels by 20 dpi. Neurofilament light chain (NfL) in the serum, used as a biomarker for neuroaxonal injury and treatment efficacy, required a longer time than 20 dpi to return to normal level. NfL is a well-established marker for neuroaxonal damages, with elevated levels reflecting the extent of neurological injury in neurodegenerative diseases (31, 32). Following a single IV+IM treatment, NfL levels remained elevated at 20 dpi, suggesting that neuroaxonal injury may have been ongoing at that time. However, considering the relatively long half-life of circulating NfL, which ranges from weeks to months (33, 34), the NfL levels detected in our study may have overstated the actual extent of neuronal lesions. After five IV treatments, NfL levels approached the normal range by 60 dpi, suggesting that neuronal damage had subsided well before the experimental endpoint. Due to ethical constraints that limit blood collection from sick animals, kinetic NfL data could not be obtained during the course of the experiments. Nevertheless, the decline in NfL levels between 20 and 60 dpi suggests that additional treatments may have contributed to reducing potential neurological damage. Along with our previous findings (24), the therapeutic efficacy of the RVC20+RVC58 mAbs cocktail in the present study was supported by a combination of clinical, laboratory and virological parameters.

Although symptomatic rabies is invariably fatal, macroscopic lesions in the CNS are mild or absent (35). The most frequent histopathological finding is infiltration of lymphocytes and microglial nodules (35, 36, 37), with inflammation generally more severe in the midbrain and medulla (35). While we detected a relative increase in the T lymphocyte population in the brains of treated animals, microglia are the main cell population that responds to RABV infection and mAb treatment. In our study, microglia exhibited slight time-dependent reactivity to RABV infection; however, this reactivity was more intense during the mAbs therapy. This treatment-dependent reactivity was also reflected in the microglial transcriptome: microglia from animals treated with RVC mAbs exhibit a distinct profile compared to those from LALA-treated or non-treated animals. On the one hand, the increased expression of interferons and interferon-stimulated genes in the brain of RVC-treated animals is consistent with increased expression of the M1-associated markers *Cxcl9* and *Nos2*, indicating a possible antiviral response (38). On the other hand, elevated brain expression of *Il10* and *Tgfb* parallels increased microglial expression of *Arg1*, *Vegfa* and *Tgm2*, classical M2 markers (39). Although microglia polarization cannot be defined as a simple M1/M2 dichotomy, the transcriptomic profile observed in RVC-treated mice is broadly consistent with a neuroprotective or reparative phenotype. Similarly, microglia in non-infectious brain conditions express *Arg1*, *Vegfa*, and *Tgm2*, a profile associated with a reparative role after ischemic stroke in a mouse model (40).

From the transcriptomics analyses, we also noticed that RABV infection induced the expression of the FcγRs *Fcgr1*, *Fcgr2b* and *Fcgr4* in microglial cells, in a similar fashion to other studies on viral infections (41, 42). The mAbs cocktail treatment did not affect the expression of *Fcgr1* in comparison to non-treated animals, but induced a higher expression of *Fcgr4*, the mouse orthologue of the human *Fcgr3a* (43). In humans, FcγRI (CD64) and FcγRIIIa (CD16a) are activating receptors regulated by Th1 cytokines such as IFN-γ (41) and involved in the pathogenesis of neurodegenerative diseases (44) and mental disorders (45). Notably, the mAbs treatment also increased the expression of *Fcgr2b* in microglia, the only known inhibitory FcγR. The expression of this receptor is induced by Th2 cytokines, such as IL-4, IL-10 or TGF-β. Although FcγRIIb (CD32b) can be co-expressed with activating FcγRs, it negatively regulates immune cell activation (46). Overexpression of this receptor can be detrimental, as in the case of Parkinson’s disease, where α-synuclein binds to FcγRIIb, inhibiting microglial phagocytosis and favoring disease progression (47). In the context of rabies therapy, however, these contrasting profiles of inhibitory and activating FcγRs appear to promote a neuroprotective environment while supporting antiviral responses. Although human mAbs may not engage mouse FcγRs with full efficacy, evidence suggests that human antibodies retain comparable FcγR-mediated biological activities in mice (48, 49), supporting the translational relevance of our findings.

We have previously demonstrated that RVC20 and RVC58 have complementary Fab and Fc properties (20, 23, 24), and our functional data presented here corroborate these differences. Structural modeling may provide partial explanation: RVC20 binds laterally to the RABV-G (21, 22) and based on published literature, the access to this binding site may be restricted depending on the topography of viral glycoproteins on the virion or infected cell surface (50, 51, 52), potentially contributing to higher IC50 of RVC20 compared to RVC58. In contrast, RVC58 binds to RABV-G in an apical position, which may allow for a denser Fc disposition on virion surface, likely enhancing FcγR engagement. Furthermore, flexibility of antibodies on the hinge regions has been associated with effector functions (53, 54), and the balance between activating and inhibitory FcγR simultaneously expressed on immune cells is important to determine the cellular response (55, 56). FcγRI binds to monomeric IgGs. However, monovalent binding is insufficient to generate a signal through FcγRIIa and FcγRIIIa. These receptors require the binding of multiple Fc fragments of antibodies present in immune complexes to induce a cellular response (46). The accessibility of multiple Fc fragments may explain why RVC58 outperforms RVC20 in binding to both FcγRIIa and FcγRIIIa when complexed with RABV. These observations suggest that beyond neutralization potency, the spatial organization of the mAbs in complex with their targets may also impact Fc-mediated effector functions and contribute to therapeutic outcome.

There have been various efforts to deliver therapeutic antibodies into the brain for symptomatic rabies treatment (7, 11). Previously, we described that the intracerebroventricular (ICV) administration of the mAb cocktail RVC20 and RVC58 cured symptomatic rabid mice (24). However, the ICV route may be unrealistic because it is highly invasive and requires advanced neurosurgical techniques that are not always available in all medical settings particularly in low- and middle-income countries where rabies is still endemic. The challenge is then to deliver these therapeutic antibodies to the CNS using more hands-on injection routes. Among alternative strategies, the most promising method for therapeutic antibodies delivery is receptor-mediated transcytosis (57). Receptor-mediated transcytosis consists of modified antibodies that target CD98 heavy chain or transferrin receptors that are heavily expressed at the blood-brain barrier. This technique can increase antibody concentrations in the brain by up to 30-fold after intravenous administration compared to non-modified antibodies. However, in this case, mAbs contain LALA modifications (58, 59, 60), which compromise Fc-mediated effector functions that are important for rabies therapy. Finally, a recent study used the cell-penetrating peptide SynB1 to increase brain delivery of a triple anti-rabies mAb cocktail, achieving only moderate success despite using three distinct antibodies. This limited efficacy may stem from inefficient effector functions that were not characterized (61).

Here we report that the unmodified anti-rabies human mAbs RVC20 and RVC58 can improve the survival rate of early symptomatic rabies in mice when primarily administered intravenously and intramuscularly. In this therapeutic setting, RVC58 monotherapy performed better than RVC20 and the RVC20+RVC58 cocktail, achieving a survival rate of 80%. Although RVC20 and RVC58 bind to different antigenic sites and do not compete in neutralizing RABV, we hypothesize that their distinct binding topography to the viral glycoprotein, and FcγR engagement in the presence of virions may partially explain why the treatment was more effective with RVC58. However, the lower total antibody dose administered in monotherapy (25 mg/kg) compared to the cocktail (50 mg/kg) may have also contributed to this outcome, possibly by eliciting a more moderate immune response, as suggested by the slightly delayed increase in clinical scores observed in monotherapies. Additionally, it should be noted that the 60-day observation period to assess the efficacy of mAb treatment was performed using a single RABV isolate in female Balb/c mice, which limits the generalizability of these findings. Therefore, further investigation is required to elucidate the precise mechanisms underlying the more efficient mAb therapies.

The WHO recommends using at least two different mAbs with non-overlapping epitopes in PEP to broaden the coverage of RABV neutralization and reduce the risk of escape mutants selection (62). While this rationale is well established for prophylaxis, the priorities for therapy against symptomatic rabies may differ, where fast viral clearance and immune response would have higher importance. In under-resourced settings, prompt access and lower cost for treatment could be determining factors, making monotherapy with an efficient antibody a potentially relevant option, especially if paired with early and accurate diagnosis. In this therapeutic context, a particular attention should be given to the binding topography of mAbs to RABV-G and their capacity to engage FcγRs and full effector functions, in addition to neutralization potency. This finding paves the way for new avenues in the development of an efficient, low-cost, and easy-to-use anti-rabies treatment.

## Materials and methods

### Ethics statement

All animal experiments were performed according to the French legislation and in compliance with the European Communities Council Directives (2010/63/UE, French Law 2013–118, February 6, 2013) and according to the regulations of Pasteur Institute Animal Care Committees. The Animal Experimentation Ethics Committee (CETEA 89) of the Institut Pasteur approved this study (160111; APAFIS#15772-2018062910157376 v5) before experiments were initiated. All animals were handled in strict accordance with good animal practice.

### Mice model for rabies

Upon arrival, female SPF Balb/cJRj mice (7 weeks-old; Janvier Laboratories) were provided with food, water, and enrichment items (cottons for nest) for the cages. The animals were randomly assigned to the experimental groups prior to infection. After one week of acclimatation period, the animals underwent isoflurane anesthesia and were inoculated with 4×10^3^ FFU (fluorescent focus units) of a pathogenic RABV (isolate 8743THA, EVAg collection, Ref-SKU: 014 V-02106, isolated from a human bitten by a dog in 1983 in Thailand) in a total volume of 100 µL (diluted in 0.85% saline, Oxoid #BO0334E), injected into the gastrocnemius muscle of both hind limbs (two injections of 25 µL in each limb). The animals were daily monitored for weight and clinical symptoms daily and scored according to the classification indicated in Appendix Table S1, which categorically scored the observations based on the appearance of animals, weight, behavior, control of movement, and the degree of symptoms. When the mice started to have locomotion difficulties, DietGel (ClearH2O #72-06-5022) was provided to all animals to ease the access to food and hydration.

### Behavioral tests

To measure forelimbs muscle strength of mice, we used the grip strength test as previously described (63) and a grip strength meter (Bioseb #BIO-GS4). The mice were tested at different time-points post-infection. Each animal was tested three times, and the grip strength, expressed in grams, represents the mean of three consecutive measures.

To assess motor coordination, we used the rotarod test (64) and an accelerating rotarod apparatus (Bioseb #BX-ROD-M), using a previously described acceleration scheme (24). The mice were tested at different time-points post-infection. Each animal was tested twice, with a 10-minute interval between each session, and the results are expressed as the mean latency time to fall off the rod. If the mouse was not able to move due to paralysis or apathy, their latency time was recorded as 0 s.

### Intravenous rabies treatment with monoclonal antibodies

At 6-, 7-, or 8-days post-infection (dpi), infected or non-infected animals received a 1:1 cocktail of the IgG1 monoclonal antibodies RVC20 and RVC58 (24) intramuscularly at a dose of 2+2 mg/kg, in a final volume of 200 µL/mouse, administered into the gastrocnemius muscle of both hind limbs (two injections of 50 µL in each limb). Afterwards, a 1:1 cocktail of the same monoclonal antibodies was intravenously administered via a single retroorbital injection at a dose of 25+25 mg/kg, in a final volume of 100 µL/mouse. The intravenous injections were performed under gas anesthesia with isoflurane (Isovet 100%, Piramal Health) and repeated 5 times, every 5 days. Infected non-treated animals received saline solution only. We also tested the LALA versions of the mAbs (RVC20-LALA and RVC58-LALA) which have two point-mutations on their Fc region (L234A and L235A) that abolish their binding affinity to FcγRs and complement. The production method of the monoclonal antibodies is described elsewhere (24).

### Kinetics of anti-rabies treatment and tissue sampling

At 6-dpi, infected or non-infected animals received one dose of 1:1 cocktail of the monoclonal antibodies RVC20 and RVC58 (named RVC) or a 1:1 cocktail of the modified monoclonal antibodies RVC20-LALA and RVC58-LALA (named LALA) via IV+IM as described above. The intravenous injections were performed under gas anesthesia with isoflurane (Isovet 100%, Piramal Health). Infected non-treated animals received saline solution only. At 8-, 10-, 12- and 20-dpi the animals were deeply anesthetized with a mixture of ketamine (Imalgène 1000, at 125 mg/kg) and xylazine (Rompun, at 5 mg/kg), blood was collected by intracardiac puncture, and the animals were perfused with PBS. The blood was stored in 1.5 mL tubes and allowed to coagulate at room temperature for at least 30 minutes. The tubes were centrifuged at 2,000 × g during 10 min at 4°C and stocked at −80°C. The brain was extracted from the skull, the two brain hemispheres were separated by a median incision: one hemisphere was fixed in 10% neutral-buffered formalin, the other hemisphere was separated in two macroscopic regions, (1) the telencephalon, composed of the cerebral cortex, the olfactory bulbs and the hippocampus, and (2) the brainstem + the cerebellum, and stocked at −80°C. The spinal cord was extracted from the vertebral column and frozen at −80°C.

### ELISA to detect human antibodies in serum and brain

Frozen tissues were briefly thawed on ice before being processed. Lysis buffer was prepared in PBS mixed with 1% NP40, protease and phosphatase inhibitors (Roche #04693159001 and #04906837001; 1 tablet of each per 10 mL) in advance. A hundred-milligram of each tissue sample was added to Lysing Matrix M tube (MP #6923100) containing 1 mL of lysis buffer and then homogenized in Bead-mill 24 homogenizer (OMNI #19523) for 20 seconds at 4 m/s, twice with 2 minutes repose between. The homogenized samples were further incubated in ice for 10 minutes and then placed in a centrifuge to spin down debris at 12,000 x g for 20 minutes. Supernatants were collected and diluted in dilution buffer (PBS mixed with 0.05% Tween 20 and 1% BSA (Sigma #A9085-25G) in 1:20 ratio. Serum samples were diluted in 1:500. Hydrophilic 384-well plates (Thermo #464718) were coated with capture antibodies (JIR #709-006-098) diluted in coating buffer (sodium bicarbonate (Sigma #C3041-50CAP) in water) to 1 μg/mL and kept at 4°C overnight. On the following day, the plates were washed with PBS containing 0.05% Tween 20 (PBST) three times and treated with 60 μL of blocking buffer (5% BSA added to PBST) to be incubated at room temperature for one hour. After washing the plates three times, 25 μL of samples and serially diluted seven standards (RVC20 or RVC58, 3-fold diluted from 300 to 0.4 ng/ml) along with a blank were loaded on plates in duplicate and kept on rotator in room temperature for two hours. Afterwards, the plates were washed, and HRP-tagged detection antibodies were added in 0.02 μL/mL final concentration for one-hour incubation on rotator, room temperature. At the end of procedure, the plates were washed again before TMB (Thermo #34028) was added for 10-minutes incubation at room temperature, followed by addition of 2N sulfuric acid to stop the reaction. The plates were read with 450 nm absorbance filter in Victor Nivo microplate reader (PerkinElmer). The concentration values are expressed in nM, taking into account the concentration in µg/mL and the estimated molecular weight of an IgG of 150 kDa.

### Neurofilament assay with serum

Before NfL dosing, the serum samples (45 µL) were treated with 1% Triton X-100 (5 µL) at room temperature for 2 h. The serum levels of NfL were measured using the commercially available single molecule array (SIMOA) assay NF-Light v2 Advantage kit (Quanterix #104073) on an HD-X analyzer following the manufacturer’s instructions.

### RNA isolation and transcriptional analyses by quantitative PCR

Total RNA from brainstem + cerebellum and telencephalon was extracted using the Direct-zol RNA MiniPrep kit (Zymo Research #R2052). Total RNA from spinal cord was extracted using the Direct-zol RNA MicroPrep kit (Zymo Research #R2062,). In both cases, the tissue fragments were weighed and added to Lysing Matrix M tubes (MP #6923100) containing 1mL of ice-cold DMEM (Dulbecco’s Modified Eagle Medium, Gibco) supplemented with 1% penicillin/streptomycin (15140148, Thermo Fisher). The tubes were then homogenized in a Bead-mill 24 homogenizer (OMNI #19523) for 20 seconds at 4 m/s, twice, with a two-minute rest period between each cycle. 125 µL of the tissue homogenate were incubated with 375 µL of Trizol LS (Invitrogen #10296028) and the extraction was performed according to the manufacturer’s instructions. RNA was reverse transcribed to first strand cDNA using the SuperScript™ IV VILO™ Master Mix (Invitrogen #11766050). qPCR was performed in a final volume of 10 μL per reaction in 384-well PCR plates using a thermocycler (QuantStudio 6 Flex, Applied Biosystems) and its related software (QuantStudio Real-time PCR System, Design and Analysis Software v. 2.7.0, Applied Biosystems). In each well, 2.5 μL of cDNA (12.5 ng) were added to 5 μL of Power SYBR Green PCR Master Mix (4367659, Applied Biosystems) and 2.5 μL of nuclease-free water containing 1 µM of primer pairs. The mouse gene targets were selected for quantifying host inflammatory mediators’ transcripts in the CNS using the *Gapdh* gene as a reference (Qiagen #249900; QuantiTect Primer Assays: *Gapdh*: QT01658692; *Ifnb*: QT00249662, *Ifng*: QT02423428, *Il6*: QT00098875, *Il10*: QT00106169; *Tgfb*: QT00249711; *Tnfa*: QT00104006). The amplification conditions were as follows: 50 °C for 2 min, 95°C for 10 min, and 45 cycles of 95 °C for 15 s and 60 °C for 1 min, followed by a melt curve from 60 °C to 95 °C. Variations in gene expression were calculated as the n-fold change in expression in the tissues from the infected mice compared with the tissues of the mock-infected group using the 2^-ΔΔCt^ method (65). To quantify the viral load in the CNS, we targeted the gene of the nucleoprotein as previously described (24).

### Immunohistochemistry

All samples were fixed in 10% neutral-buffered formalin for 48 hours and embedded in paraffin. Sections were cut to 4 μm thickness and processed for routine histology using hematoxylin and eosin staining and chromogenic Immunohistochemistry with hematoxylin counterstaining. IHC analysis was performed using the primary antibodies rabbit anti-Iba-1 (Wako #019-19741; dilution 1:1000) to detect microglia, and with a rabbit anti-P49-1 antibody (in house; dilution 1:1000) (36) to detect RABV. We also used a rabbit anti-CD3 (Dako #A0452; dilution 1:400) to detect T lymphocytes, and with a rat anti-B220 (CD45R) antibody (BD Pharmigen #550286; dilution 1:500) to detect B lymphocytes. The secondary antibodies were a biotinylated anti-rabbit (Dako #E0432; dilution 1/600) and a biotinylated anti-rat (Vector Laboratories #BA-9400; dilution 1:600). All slides were scanned using the AxioScan Z1 (Zeiss) system, and images were visualized with the Zen 3.9 software. Cortex images obtained from anti-Iba1 staining were converted into 8-bit JPG files using FIJI and then overlaid with a threshold-set mask to measure the antibody-stained area occupied by resting / reactive microglia. We also quantified T and B lymphocytes staining using the QuPath software (version 0.5.1) (66). Briefly, we defined regions of interest (ROIs) by drawing a contour delimitating the whole area of interest (cerebral cortex, thalamus, midbrain, cerebellum, pons and medulla oblongata). The area of CD3 (T lymphocytes) and B220 (B lymphocytes) positive staining was calculated within the ROIs, and the results were expressed as percentage of positive stained area.

### Microglia isolation, bulk sequencing and transcriptome processing

Microglia from mice brain were isolated using Adult Brain Dissociation Kit, according to the manufacturer’s protocol (Miltenyi Biotec #130-107-677). A brief procedure was the following: On 9 dpi, antibody-treated and non-treated animals were sacrificed with CO_2_. Brains were removed and washed with ice-cold D-PBS immediately and cut into small pieces to aid enzyme digestion. The tissues were mechanically dissociated with enzymes and then placed on a rotator at 37°C for 5 minutes. After the digestion, the tissues were gently dissociated again with pipettes and centrifuged at 300 x g for 10 minutes. Live cells were separated from tissue debris and red blood cells using respective solutions and multiple steps of centrifuge and then labelled with anti-CD11b-APC (Miltenyi Biotec #130-113-793) and anti-CD45-FITC (Miltenyi Biotec#130-116-500), which are classical markers used to isolate microglia. Up to this point, all tissues / cells were manipulated in ice-cold condition. Afterwards, labelled cells were fixed with 4% paraformaldehyde for 30 minutes and then resuspended in D-PBS for FACS sorting (BD Biociences). Double-labelled microglia were isolated through multiple gatings of size, complexity and fluorescence, and then subjected to RNA extraction.

Total RNA from fixed microglia was extracted using RecoverAll™ Total Nucleic Acid Isolation Kit (Invitrogen #AM1975) and Trizol LS (Invitrogen #10296010) using a slightly modified version of manufacturers’ protocols. In brief, microglia pellets were suspended in mixture of protease and RNAsin (Promega #N2611) and incubated at 50°C for 15 minutes followed by another 15 minutes at 80°C. Isolation Additive solution was added to the lysate and mixed before being suspended in appropriate volume of Trizol LS. After extracting RNA according to the manufacturer’s protocol, the RNA pellets were treated with DNase I (Promega #M6101) and then subjected to second Trizol LS extraction. Collected RNA samples went through ethanol precipitation as previously described (67) to increase the purity of RNA.

Library preparation was carried out using the Illumina Stranded mRNA Library Preparation Kit (Illumina, USA) according to manufacturer’s protocol, with modifications based on the quality control assessment of the samples. Sample integrity and RNA quantity were evaluated using RNA 6000 Pico Kit (Agilent Technologies, USA) on Agilent Bioanalyzer. Sequencing was conducted on a NextSeq2000 platform (Illumina, USA), generating on average 32M reads (115-base single-end reads) per sample.

The RNA-seq analysis was performed with Sequana 0.18.0 (68). We used the RNA-seq pipeline 0.20.0 (https://github.com/sequana/sequana_rnaseq) built on top of Snakemake 7.32.4 (69). Briefly, reads were trimmed using fastp 0.23.2 (70) then mapped to the *Mus musculus* GRCm39 genome assembly from Ensembl using STAR (71). FeatureCounts 2.0.1 (72) was used to produce the count matrix, assigning reads to features using corresponding annotation v109 from Ensembl with strand-specificity information. Quality control statistics were summarized using MultiQC 1.16 (73). Statistical analysis on the count matrix was performed to identify differentially regulated genes. Clustering of transcriptomic profiles were assessed using a Principal Component Analysis (PCA). Differential expression testing was conducted using DESeq2 1.38.3 library (74) scripts indicating the significance (Benjamini-Hochberg adjusted p-values, false discovery rate FDR < 0.05) and the effect size (fold-change) for each comparison.

### Immune response-associated gene expression and enrichment analysis

Differentially expressed genes (DEGs) were all obtained from comparing infected sets to non-infected (NI) set. Each set of DEGs (NT vs. NI, RVC vs. NI, or LALA vs. NI) was respectively described as NT, RVC or LALA throughout this article to simplify discourse. Overlapping between or exclusive genes to the compared DEG sets were counted using ggVennDiagram package within R studio. To present gene ontologies (GOs) related to immune responses, first enrichment analysis was performed for each DEG sets using mouse-specific annotation package org.Mm.eg.db (Bioconductor), and then the GO terms in Biological Process were filtered through a collection of keywords related to immunity as previously described (67). geneRatio was calculated from dividing the number of DEGs by the number of genes in each ontology set. Panels of GO terms were prepared using ggplot package, and the length of terms in y axis was limited to 60 characters for display. DEGs from immune response-associated GO terms and common to all three DEG sets were sorted to the differences of log2 fold-changes, standardized to Z-score for each DEG, and presented as heatmaps (aheatmap in NMF package).

### Rabies virus neutralization assay

RABV neutralization by RVC20 and RVC58, alone or in cocktail, was evaluated using a modified rapid fluorescent focus inhibition test (RFFIT), recommended as the reference technique by WHO (75). We tested the neutralization of these mAbs against the RABV strain used in this study (isolate 8743THA), and against three other Lyssaviruses: the cell culture-adapted rabies challenge virus (CVS-11), a vampire bat-related RABV (isolate 9001FRA), and the European bat lyssavirus 1 (isolate 8918FRA).

Briefly, a constant dose of virus, determined by a previous titration to give a percentage of cell infection between 80% and 95%, was incubated with a three-fold serial dilution of each mAb. After 24h of incubation (CVS-11 and the isolate 9001FRA) or 48h of incubation (isolates 8743THA and 8918FRA) (20) at 37°C in a humid atmosphere under 5% CO_2_, the cell monolayer was fixed with 80% (v/v) acetone and labelled with a fluoresceinated anti-rabies nucleocapsid antibody (Merck #5100). The IC50 was calculated by calculating the concentration of mAbs resulting in 50% of virus neutralization.

### Rabies virus isolation from mouse spinal cords

We added 50 µL of the spinal cord homogenate (prepared as for the RNA extraction) to 4×10^4^ murine neuroblastoma cells (ATCC) in a Lab-Tek™ Chamber Slide System (Nunc #154534), in a final volume of 400 µL of DMEM supplemented with 10% fetal calf serum. To determine the assay’s limit of detection, we also prepared eight serial 3-fold dilutions (from 3,000 to 1.4 fluorescent focus-forming units (FFU)/50 µL) of the titrated RABV stock (isolate 8743THA) that was used to infect the animals. After 72 hours of incubation at 37 °C, the cells were fixed and permeabilized with 80% acetone. Then, the cells were stained with a FITC-conjugated anti-rabies monoclonal globulin (Fujirebio #800-092) and observed with the EVOS M5000 microscope (Invitrogen).

### RVC20 and RVC58 binding Fcγ receptors

We tested the RVC20 and RVC58 binding to human FcγRs using the Lumit® FcγR binding immunoassays: FcγRI (Promega #W7080), FcγRIIa H131 (Promega #W7070), FcγRIIIa V158 (Promega #W7050), and FcγRIIb (Promega #W7030). We initially tested the binding capacity of the mAbs to FcγRs using 4-fold dilutions of RVC20 only, RVC58 only, or RVC20+RVC58 cocktail (from 2.5×10^2^ to 6×10^-5^ µg/mL) in 384-wells plates following the manufacturer’s instructions. Luminescence was measured with the Victor Nivo plate reader (Perkin Elmer).

We also tested the FcγRs binding capacity of the mAbs after neutralizing RABV. To this aim, we incubated 10-fold dilutions of RVC20 only, RVC58 only, or RVC20+RVC58 cocktail (from 1×10^3^ to 1×10^-4^ µg/mL) with 1×10^5^ FFU of the pathogenic RABV (isolate 8743THA) during 1h at 37°C in 96-wells plates. Afterwards, we assessed the binding of the mAb-RABV complex to FcγRI (Promega #W7080), FcγRIIa H131 (Promega #W7070), FcγRIIIa V158 (Promega #W7050), and FcγRIIb (Promega #W7030) following the manufacturer’s instructions. Luminescence was measured with the Glomax Multi reader (Promega).

#### Modelling the RVC20 and RVC58 interaction with the RABV-G

The RVC20 complex was generated by superimposing the available structure of the mAb’s single-chain variable fragment (scFv) bound to RABV G domain III (PDB: 6TOU) (21) on domain III in the full ectodomain of the RABV G trimer. For this purpose, we took the atomic model of the available structure of G in complex with Fab RVA122 (PDB: 7U9G) (22). The alignment resulted in a root mean square deviation (RMSD) of 0.685Å for 61 pairs of matching Cα atoms of G domain III. We then superimposed the atomic model of a full-length immunoglobulin G (IgG) molecule available in the database (PDB: 1IGT) (76) on the variable domains of the heavy and light chains of RVC20 bound to the G trimer (RMSD = 0.963Å for 195 matching pairs of Cα atoms in both heavy and light chains). For RVC58, we showed previously that it shares epitope with RVA122 (23), which was crystallized in complex with RABV G (PDB: 7U9G) (22). We therefore used this structure to superimpose the same full-length IgG molecule on the variable domains of the RVA122 heavy and light chains, which resulted in an RMSD of 1.305 for193 Å for 193 matching pairs of Cα atoms in both heavy and light chains.

#### Statistics

Statistical analysis was performed using Prism software (GraphPad, version 10.4.1, San Diego, USA). Quantitative data were compared across multiple groups using the Kruskal-Wallis test followed by the Dunn’s multiple comparisons test. The Unpaired T test was used to compare two groups. Survival curves were compared using the Log-rank (Mantel–Cox) test. If not indicated otherwise, data are expressed as the median and interquartile range. A p-value of <0.05 was considered significant, and the exact p-values are shown in the source data tables. *In vivo* data collection and analysis were not blinded due to experimental conditions. *Ex vivo* analysis was blinded (coded samples). No data were excluded from the analyses.

## Author contribution

Seonhee Kim: Conceptualization; Formal Analysis; Investigation; Visualization; Writing – original draft

Anthony Coleon: Investigation

Florence Larrous: Investigation; Resources

Lauriane Kergoat: Investigation

Said Mougari: Investigation

Julien Lannoy: Investigation

Marta Tiago: Investigation

David Hardy: Formal Analysis; Resources

Etienne Kornobis: Formal Analysis; Resources

Felix A. Rey: Methodology; Resources; Writing – review & editing

Fabio Benigni: Resources; Writing – review & editing

Davide Corti: Resources; Writing – review & editing

Hervé Bourhy: Conceptualization; Resources; Supervision; Writing – review & editing

Guilherme Dias de Melo: Conceptualization; Investigation; Formal Analysis; Supervision; Writing – original draft; Writing – review & editing

## Acknowledgements

We thank Magali Tichit; David Hing and Sarra Loulizi, Histopathology Core Facility, Institut Pasteur, for their help with histology. We acknowledge Elodie Turc for library preparation and sequencing, Biomics Platform, C2RT, Institut Pasteur, Paris, France, supported by France Génomique (ANR-10-INBS-09) and IBISA. We also thank Esma Karkeni, Single Cell Biomarkers Unit of Technology and Service (scBiomarkers UTechS), Institut Pasteur, for support with the SIMOA assays.

## Data availability

The RNA-seq data generated in this study have been deposited in the Array Express database under accession code E-MTAB-16546 [https://www.ebi.ac.uk/biostudies/arrayexpress]. The immunohistology images generated in this study have been deposited in the BioImages database under accession codes S-BIAD3374 and S-BIAD3381 [https://www.ebi.ac.uk/biostudies/bioimages]

---

**Figure EV1.**
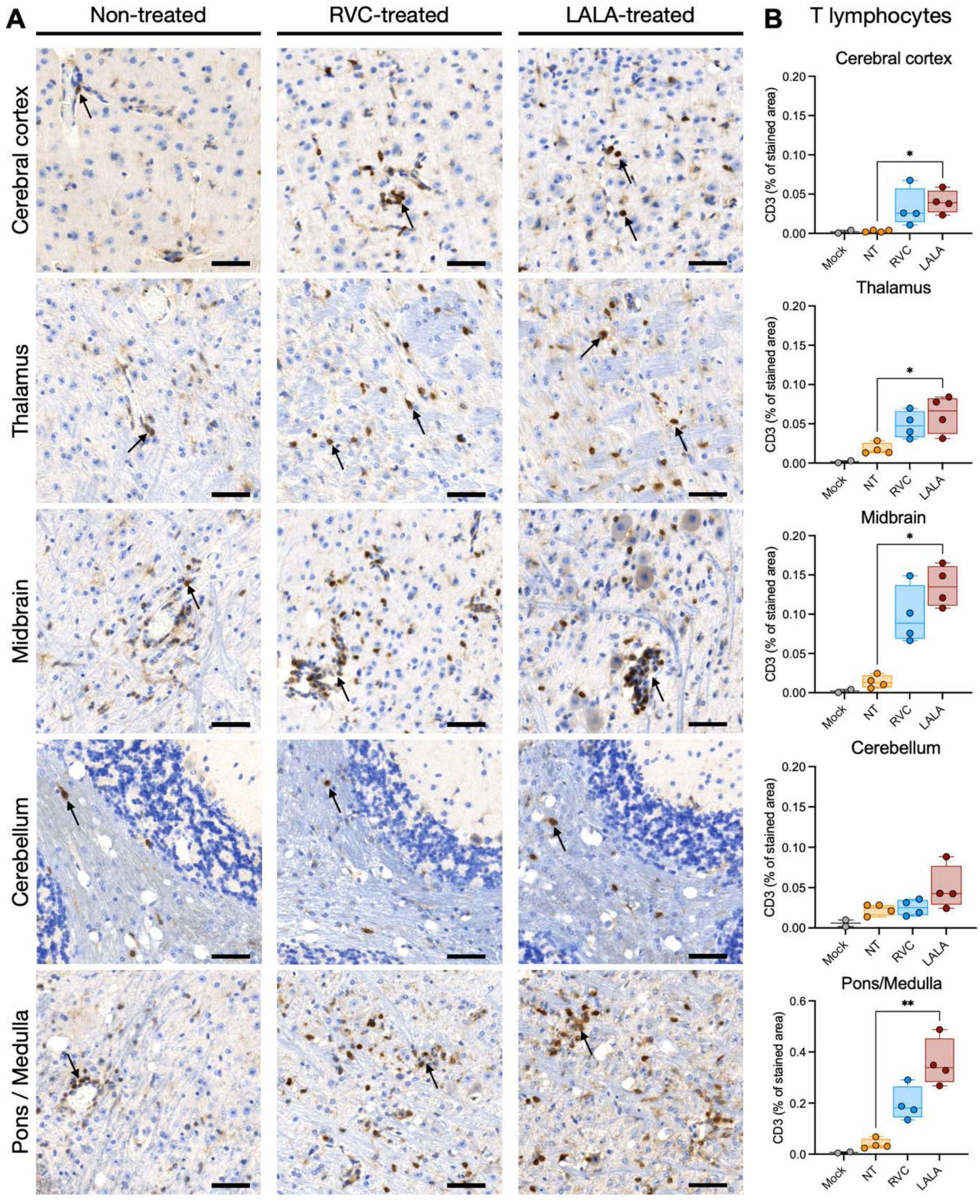
T lymphocytes in the brain of mice with symptomatic rabies. **A.** Immunohistochemical analysis showing staining for T lymphocytes (CD3, cells marked in brown, arrows) in the cerebral cortex, thalamus, midbrain, cerebellum, pons and medulla oblongata of mice under different treatments at 12 days post-infection. Scale bar = 50 µm. All images are in the same magnification. Representative images (n=4/group). **B.** Quantification of T lymphocytes in the brain of mice under different treatments at different days post-infection. Box and whisker plots (median, first and third quartiles, minimum and maximum). Individual values are also shown. Kruskal-Wallis test followed by the Dunn’s multiple comparisons test: black asterisks indicate comparisons among treatments (*p<0.05, **p<0.01). Mock (n=2); Non-treated (NT), RVC and LALA (n=4).

**Figure EV2.**
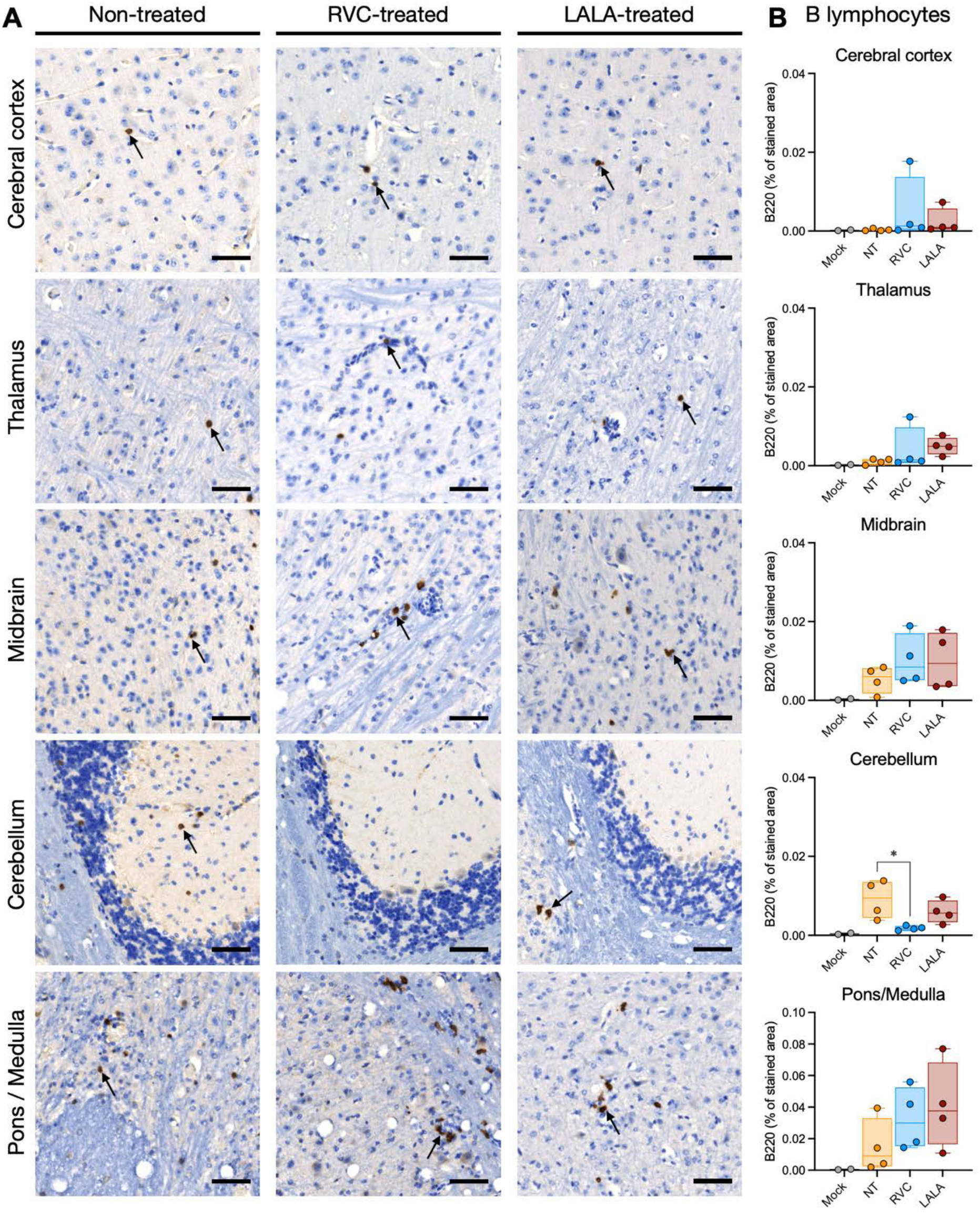
B lymphocytes in the brain of mice with symptomatic rabies. **A.** Immunohistochemical analysis showing staining for B lymphocytes (B220, cells marked in brown, arrows) in the cerebral cortex, thalamus, midbrain, cerebellum, pons and medulla oblongata of mice under different treatments at 12 days post-infection. Scale bar = 50 µm. All images are in the same magnification. Representative images (n=4/group). **B.** Quantification of B lymphocytes in the brain of mice under different treatments at different days post-infection. Box and whisker plots (median, first and third quartiles, minimum and maximum). Individual values are also shown. Kruskal-Wallis test followed by the Dunn’s multiple comparisons test: black asterisks indicate comparisons among treatments (*p<0.05). Mock (n=2); Non-treated (NT), RVC and LALA (n=4).

**Figure EV3.**
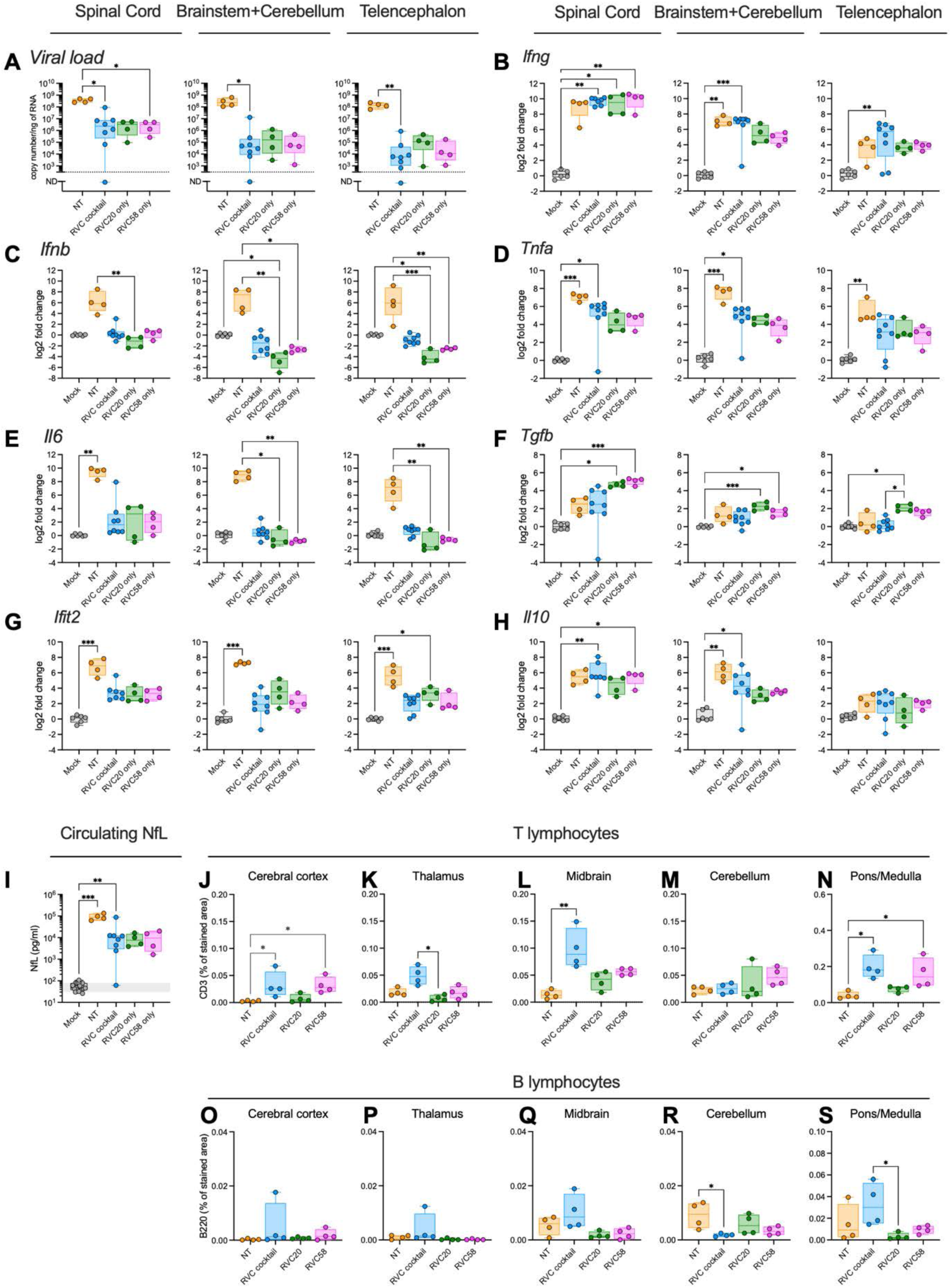
Virological and immune profile and lymphocytes infiltration in the brain of mice treated with RVC20 or RVC58 at 12 days post-infection. **A.** Rabies virus load in the spinal cord, in the brainstem + cerebellum and in the telencephalon. **B-H.** Gene expression of interferon gamma (B), interferon beta (C), tumoral necrosis factor alpha (D), interleukin 6 (E), transforming growth factor beta (F), interferon Induced protein with tetratricopeptide repeats 2 (G), and interleukin 10 (H) and in the spinal cord, brainstem + cerebellum, and telencephalon of mice under different treatments. **I.** Neurofilament light chain (NfL) levels in the serum of mice under different treatments at 12 days post-infection. **J-N.** Quantification of T lymphocytes (CD3% stained area) in the cerebral cortex (J), thalamus (K), midbrain (L), cerebellum (M), pons and medulla oblongata (N) of mice under different treatments at 12 days post-infection. **O-S.** Quantification of B lymphocytes (B220% stained area) in the cerebral cortex (O), thalamus (P), midbrain (Q), cerebellum (R), pons and medulla oblongata (S) of mice under different treatments at 12 days post-infection. Mock, NT and RVC cocktail already shown in Figures 2-3 and Figures EV3-4. RVC20 only and RVC58 only (n=4/group). Box and whisker plots (median, first and third quartiles, minimum and maximum). Individual values are also shown. Kruskal-Wallis test followed by the Dunn’s multiple comparisons test: black asterisks indicate comparisons among treatments (*p<0.05, **p<0.01, *** p<0.001, **** p<0.0001).

**Figure EV4.**
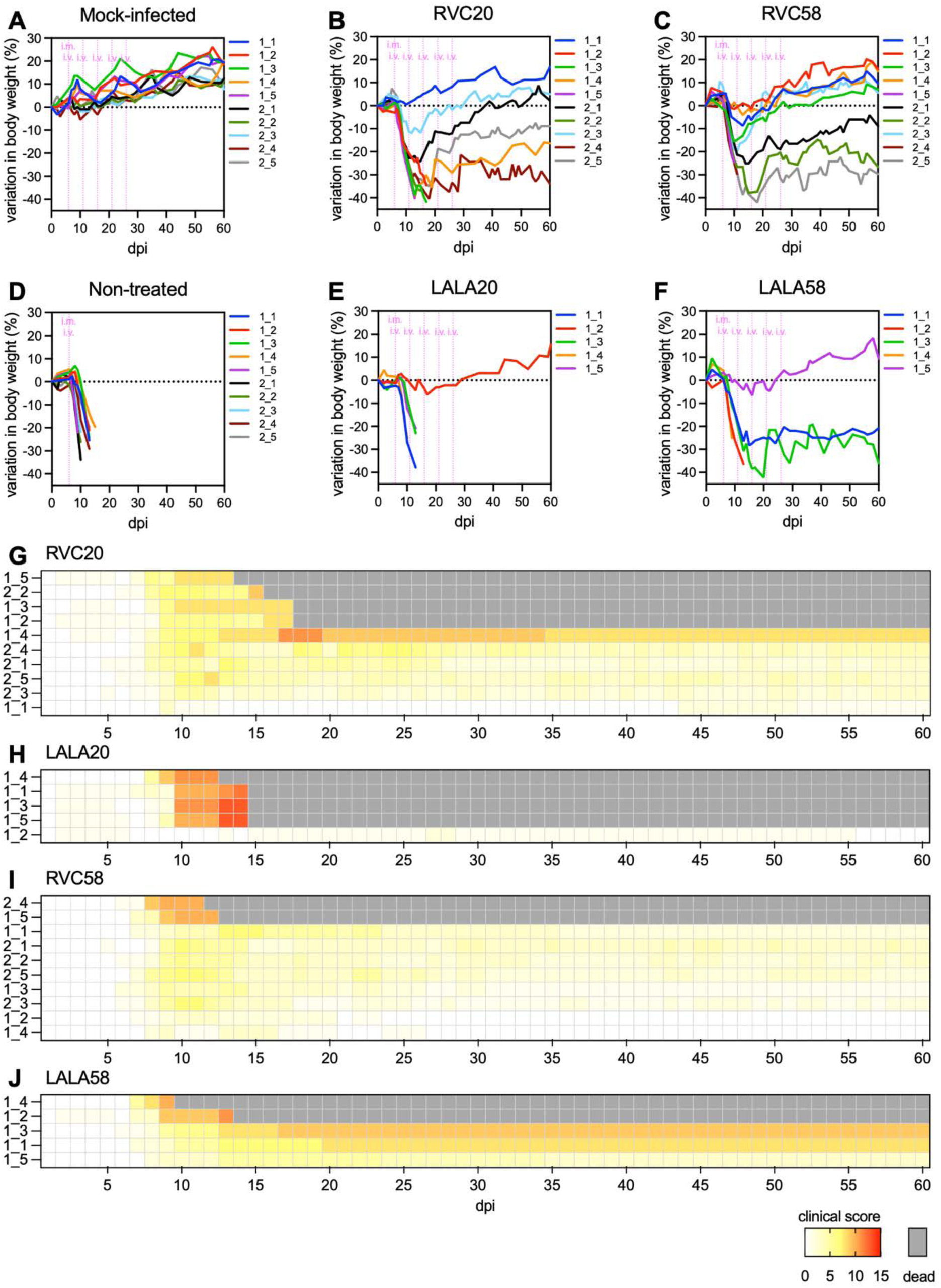
Clinical profile of rabid mice after treatment with single RVC20, LALA20, RVC58 or LALA58 monoclonal antibodies starting at 6 days post-infection. **A-F.** Individual body weight follow-up. **G-J.** Individual clinical score follow-up. Mock, NT, RVC20 and RVC58 (n=10, 2 independent experiments). LALA20 and LALA58 (n=5, 1 experiment). Data information: **G-J:** The mouse ID is as follows: The first number indicates the replicate, and the second number indicates the mouse within the replicate. For example, 1_3 indicates replicate 1 and mouse 3.

**Figure EV5.**
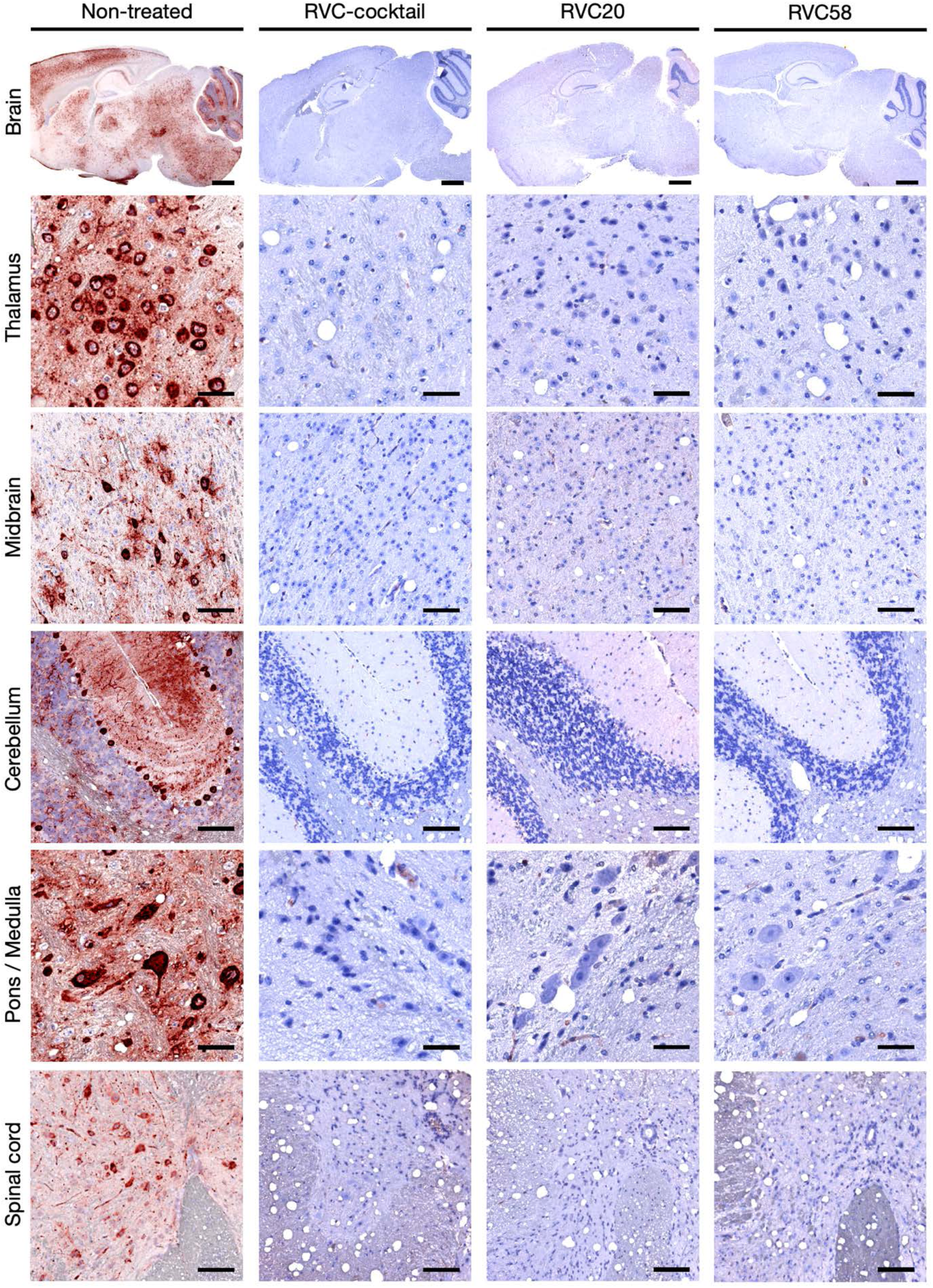
Rabies virus staining in the central nervous system of mice that survived after treatment. Immunohistochemical analysis showing staining for the rabies virus (RABV, red spots) in the whole brain, thalamus, midbrain, cerebellum, pons / medulla oblongata, and spinal cord of NT mice that died, or of mice that survived after treatment (60 dpi). Representative images. Scale bars: submacroscopic view of the brain = 1 mm; thalamus, pons / medulla oblongata = 50 µm; midbrain, cerebellum, spinal cord = 100 µm. Data information: the thalamus, midbrain, cerebellum, and pons/medulla oblongata are high-magnification views of the submacroscopic brain images in each column.

