## Appendix for "Monoclonal antibodies targeting the rabies virus glycoprotein promotes viral clearance and Fc-dependent neuroprotection in the infected brain"

Table of contents

Figures

- Appendix Figure S1. Modulated immune response in the brain under RVC20 and RVC58 monoclonal antibody cocktail treatment.
- Appendix Figure S2. Immunohistochemical analysis showing staining for the rabies virus (RABV, red spots) and for microglia (Iba1, brown cells) in the thalamus, midbrain, cerebellum, pons and medulla oblongata of mice under different treatments at different days post-infection.
- Appendix Figure S3. Viral isolation from spinal cords of animals infected with rabies virus.

Tables

- Appendix Table S1. Clinical score chart.

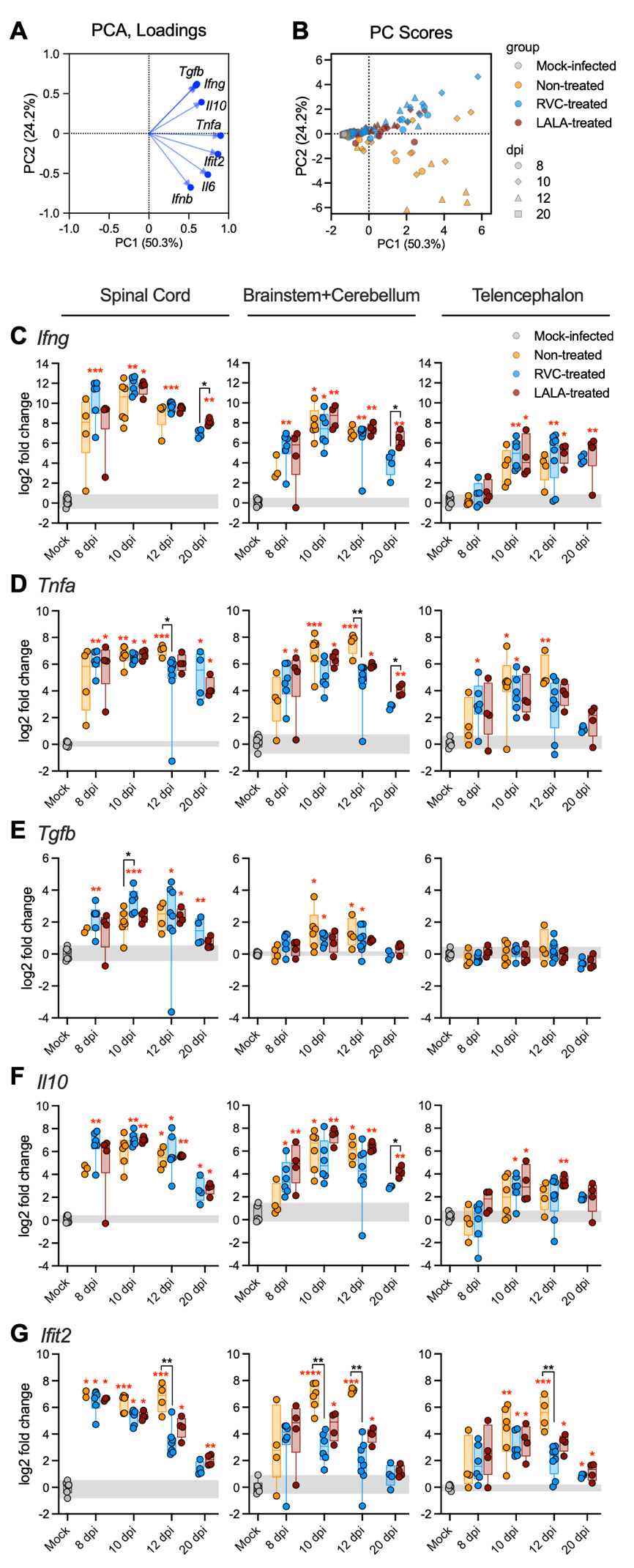
**Appendix Figure S1.** **Modulated immune response in the brain under RVC20 and RVC58 monoclonal antibody cocktail treatment. A-B.** Principal component analysis (PCA) of cytokines in the brain of mice under different treatments. (A) Variable correlation plot showing the correlation of the gene expression of different cytokines at 8, 10, 12 and 20 days post-infection. The two-first principal components explained 76% of sample variability. (B) PCA plots. Each symbol represents one animal, colored according to the treatment. **C-F.** Time-dependent gene expression of interferon gamma (C), tumoral necrosis factor alpha (D), transforming growth factor beta (E), interleukin 10 (F) and interferon induced protein with tetratricopeptide repeats 2 (G) in the spinal cord, brainstem + cerebellum, and telencephalon of mice under different treatments. Box and whisker plots (median, first and third quartiles, minimum and maximum). Individual values are also shown. The gray zone corresponds to the 95% CI of the median from the mock-infected mice. Kruskal-Wallis test followed by the Dunn’s multiple comparisons test: red asterisks indicate comparisons with the mock-infected group, whereas black asterisks indicate comparisons among treatments. Unpaired T test to compare RVC and LALA at 20 dpi (*p<0.05, **p<0.01, *** p<0.001, **** p<0.0001). Mock (n=6); Non-treated (n=4 for 8 dpi; n=6 for 10 dpi; n=4 for 12 dpi); RVC-treated (n=6 for 8 and 10 dpi; n=8 for 12 dpi; n=4 for 20 dpi); LALA-treated (n=4/time-point).

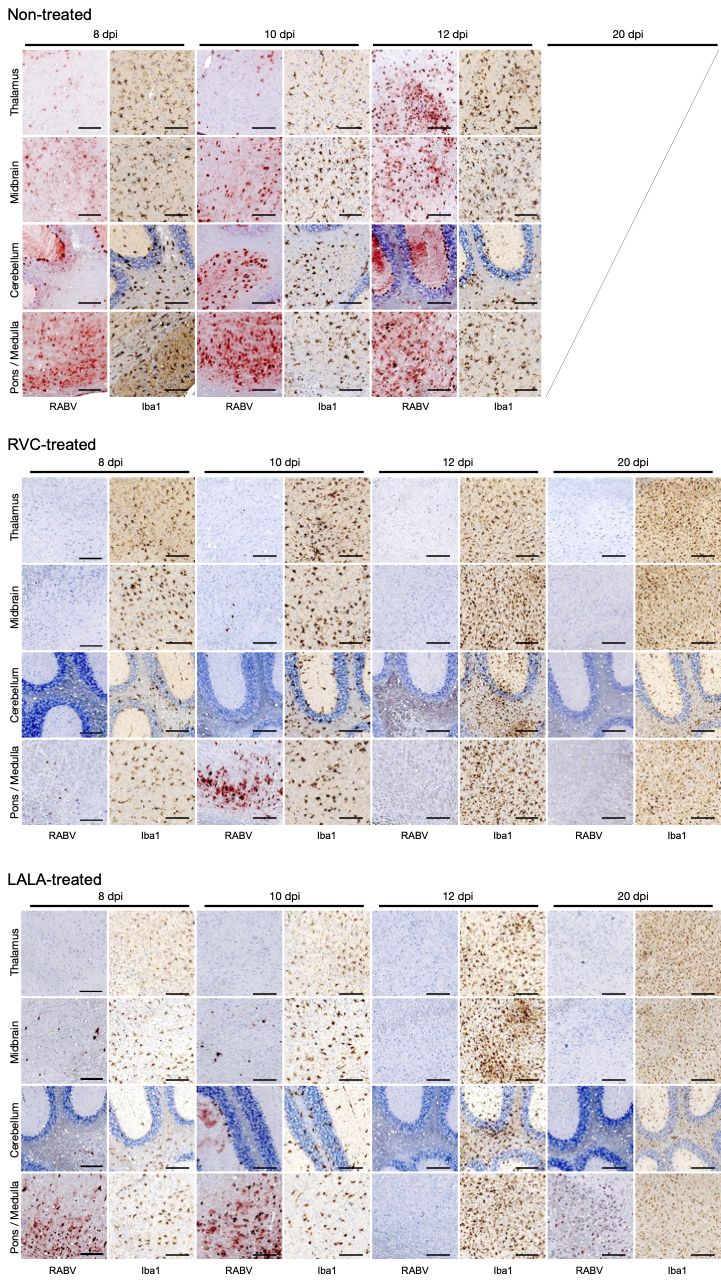

**Appendix Figure S2. Immunohistochemical analysis showing staining for the rabies virus (RABV, red spots) and for microglia (Iba1, brown cells) in the thalamus, midbrain, cerebellum, pons and medulla oblongata of mice under different treatments at different days post-infection.** Scale bar = 200 µm. Representative images. Non-treated (n=4 for 8 dpi; n=6 for 10 dpi; n=2 for 12 dpi); RVC (n=6 for 8 and 10 dpi; n=8 for 12 dpi; n=4 for 20 dpi); LALA (n=4/time-point).

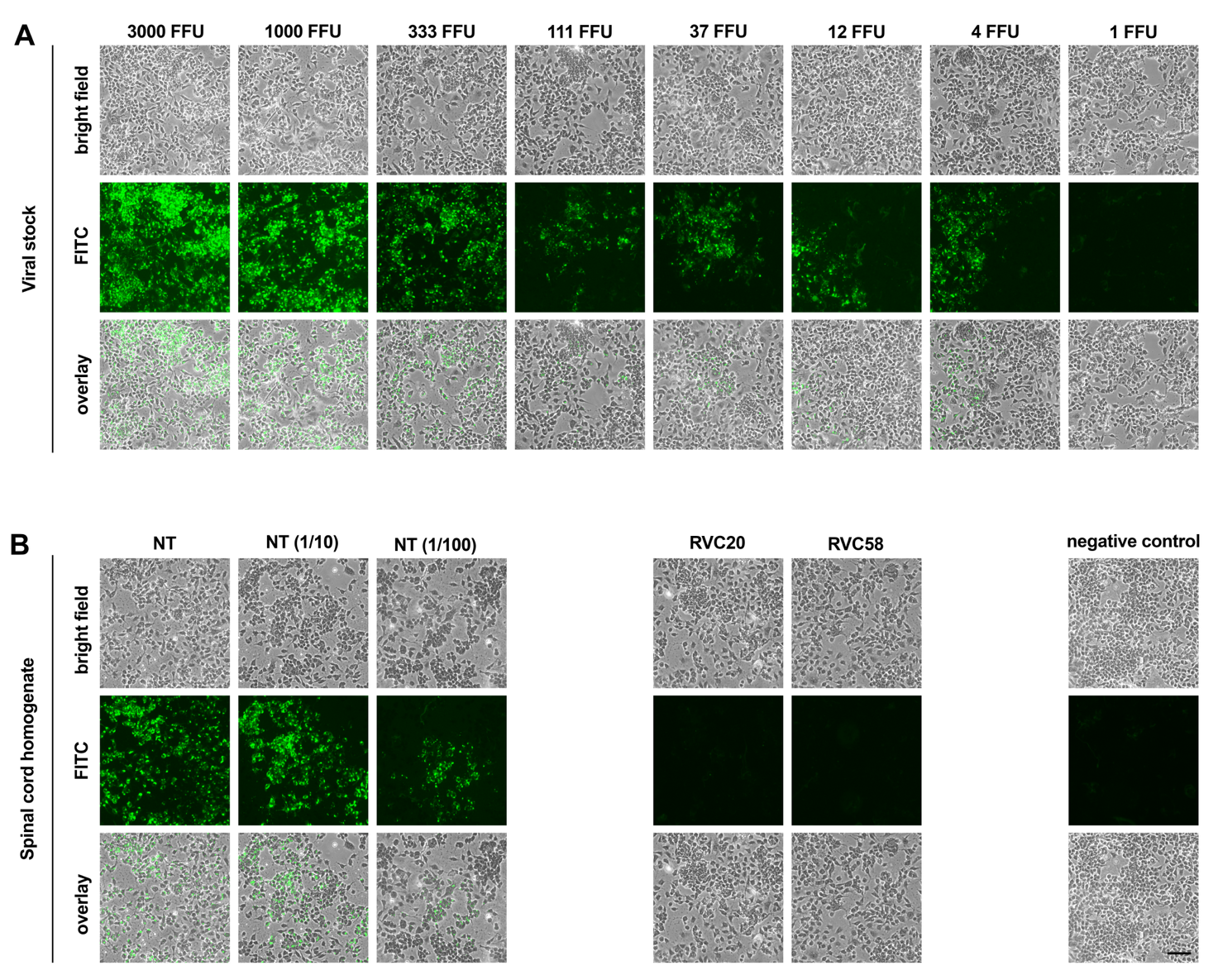

**Appendix Figure S3. Viral isolation from spinal cords of animals infected with rabies virus.** **A.** Dilution curve of the viral stock of the RABV isolate 8743THA used to infect the mice. In this setting, the detection limit is four fluorescent focus-forming units (FFU). **B.** Viral isolation from mouse spinal cords. NT: non-treated animal #2_1 (Figure 1), which died at 10 days post-infection (dpi). The spinal cord homogenate was tested undiluted (64.2 mg of tissue/mL), or diluted 1/10 and 1/100. RVC20: RVC20-treated animal #2_4 (Figure 6 and EV4), which survived until 60 dpi; undiluted spinal cord homogenate (85.8 mg of tissue/mL). RVC58: RVC58 treated-animal #2_5 (Figure 6 and EV4), which survived until 60 dpi; undiluted spinal cord homogenate (112.2 mg of tissue/mL). Scale bar = 100 µm. Representative images of the wells.

**Appendix Table S1. Clinical score chart.**

| **Parameter** | **Description** | **Score** |
| --- | --- | --- |
| **Grooming** | Normal grooming | 0 |
|  | Clear lack of grooming | 1 |
| **Locomotion** | No apparent changes | 0 |
|  | Slow movement | 1 |
|  | Overexcitation (moving too fast) and/or Disorientation (circling) | 1 |
|  | Apathy | 2 |
| **Muscle control** | No apparent changes | 0 |
|  | Asymmetric position of hindlimb when lifted | 1 |
|  | No standing on hindlimbs to explore | 2 |
|  | Slow retrieval of hindlimb while moving (unsteady movement) | 2 |
| **Involuntary movement** | No apparent changes | 0 |
|  | Tremor while sitting / moving | 1 |
|  | Whole-body convulsion when lifted | 2 |
| **Facial features** | Active and alert | 0 |
|  | Eyelids narrowed or closed but whiskers responsive | 1 |
|  | No whisker response to stimulation | 2 |
| **Paralysis** | No paralysis | 0 |
|  | Paralysis on one hindlimb | 1 |
|  | Paralysis on both hindlimbs / below lower spine | 2 |
|  | Paralysis below-neck | 3 |
|  | Full-body paralysis | 4 |
| **Weight** | No weight loss or gained compared to initial weight | 0 |
|  | Lost weight compared to the initial weight | 1 |
| **Death** | Alive | 0 |
|  | Euthanasia (humane end point) | 30 |
